# Finding Driver Mutations in Cancer: Elucidating the Role of Background Mutational Processes

**DOI:** 10.1101/354506

**Authors:** Anna-Leigh Brown, Minghui Li, Alexander Goncearenco, Anna R. Panchenko

## Abstract

Identifying driver mutations in cancer is notoriously difficult. To date, recurrence of a mutation in patients remains one of the most reliable markers of mutation driver status. However, some mutations are more likely to occur than others due to differences in background mutation rates arising from various forms of infidelity of DNA replication and repair machinery, endogenous, and exogenous mutagens.

We calculated nucleotide and codon mutability to study the contribution of background processes in shaping the observed mutational spectrum in cancer. We developed and tested probabilistic pan-cancer and cancer-specific models that adjust the number of mutation recurrences in patients by background mutability in order to find mutations which may be under selection in cancer.

We showed that mutations with higher mutability values had higher observed recurrence frequency, especially in tumor suppressor genes. This trend was prominent for nonsense and silent mutations or mutations with neutral functional impact. In oncogenes, however, highly recurring mutations were characterized by relatively low mutability, resulting in an inversed U-shaped trend. Mutations not yet observed in any tumor had relatively low mutability values, indicating that background mutability might limit mutation occurrence.

We compiled a dataset of missense mutations from 58 genes with experimentally validated functional and transforming impacts from various studies. We found that mutability of driver mutations was lower than that of passengers and consequently adjusting mutation recurrence frequency by mutability significantly improved ranking of mutations and driver mutation prediction. Even though no training on existing data was involved, our approach performed similarly or better to the state-of-the-art methods.

**Availability:** https://www.ncbi.nlm.nih.gov/research/mutagene/gene

**Author Summary:** Cancer development and progression is associated with accumulation of mutations. However, only a small fraction of mutations identified in a patient is responsible for cellular transformations leading to cancer. These so-called drivers characterize molecular profiles of tumors and could be helpful in predicting clinical outcomes for the patients. One of the major problems in cancer research is prioritizing mutations. Recurrence of a mutation in patients remains one of the most reliable markers of its driver status. However, DNA damage and repair processes do not affect the genome uniformly, and some mutations are more likely to occur than others. Moreover, mutational probability (mutability) varies with the cancer type. We developed models that adjust the number of mutation recurrences in patients by cancer-type specific background mutability in order to prioritize cancer mutations. Using a comprehensive experimental dataset, we found that mutability of driver mutations was lower than that of passengers, and consequently adjusting mutation recurrence frequency by mutability significantly improved ranking of mutations and driver mutation prediction.

## Introduction

Cancer is driven by changes at the nucleotide, gene, chromatin, and cellular levels. Somatic cells may rapidly acquire mutations, one or two orders of magnitude faster than germline cells [1]. The majority of these mutations are largely neutral (passenger mutations) in comparison to a few driver mutations that give cells the selective advantage leading to their proliferation [2]. Such a binary driver-passenger model can be adjusted by taking into account additive pleiotropic effect of mutations [3, 4]. Mutations might have different functional consequences in various cancer types and patients, they can lead to activation or deactivation of proteins and dysregulation of a variety of cellular processes. This gives rise to high mutational, biochemical, and histological intra- and inter-tumor heterogeneity that may explain the resistance to therapies and complicates the identification of driving events in cancer [5, 6].

Point DNA mutations can arise from various forms of infidelity of DNA replication and repair machinery, endogenous, and exogenous mutagens [6–9]. There is an interplay between processes leading to DNA damage and those maintaining genome integrity. The resulting mutation rate can vary throughout the genome by more than two orders of magnitude [10, 11] due to many factors operating on local and global scales [12–14]. Many studies support point mutation rate dependence on the local DNA sequence context for various types of germline and somatic mutations [9, 11, 13, 15]. For both germline and somatic mutations, local DNA sequence context has been identified as a dominant factor explaining the largest proportion of mutation rate variation [10, 16]. Additionally, differences in mutational burden between cancer types suggest tissue type and mutagen exposure as important confounding factors contributing to tumor heterogeneity.

Assessing background mutation rate is crucial for identifying significantly mutated genes [17, 18], sub-gene regions [19, 20], mutational hotspots [21, 22], or prioritizing mutations [23]. This is especially important considering that the functional impact of the majority of changes observed in cancer is poorly understood, in particular for rarely mutated genes [24]. Despite this need, there is a persistent lack of quantitative information on per-nucleotide and per-codon background rates in various cancer types and tissues.

There are many computational methods that aim to detect driver genes and fewer methods trying to rank mutations with respect to their potential carcinogenicity. As many new approaches to address this issue have been developed [25] [26], it still remains an extremely difficult task. As a consequence, many driver mutations, especially in oncogenes, are not annotated as high impact or disease related [27] even though cancer mutations harbor the largest proportion of harmful variants [28].

In this study we utilize probabilistic models that estimate background mutability per nucleotide or codon substitution to rank mutations and help distinguish driver from passenger mutations. The mutability concept has been used in many evolutionary and cancer studies (although it has been estimated in different ways). Mutability is defined as a probability to obtain a nucleotide or codon substitution purely from the underlying background processes of mutagenesis and repair that are devoid of cancer selection component affecting a specific genomic (or protein) site. The mutability is calculated using background models (mutational profiles) that are constructed under the assumption that vast majority of cancer context-dependent mutations have neutral effects, while only a small number of these mutations in specific sites are under positive or negative selection. To assure this, we removed all recurrent mutations as these mutations might be under selection in cancer. Mutational profiles are calculated by sampling the frequency data on types of mutations and their trinucleotide (for nucleotide mutations) and pentanucleotide (for codon substitutions) contexts regardless of their genomic locations. These models in the forms of mutational profiles can be used to estimate the expected mutation rate in a given genomic site as a result of different local or long-range context-dependent mutational processes.

In this paper we try to decipher the contribution of background DNA mutability in the observed mutational spectrum in cancer for missense, nonsense, and silent mutations. We compiled a set of cancer driver and neutral missense mutations with experimentally validated impacts collected from multiple studies and used this set to verify our approach and compare it with other existing methods. Our approach has been implemented online as part of the MutaGene web-server and as a stand-alone Python package: https://www.ncbi.nlm.nih.gov/research/mutagene/gene.

## Results

### Mutations not observed in cancer patients have low mutability

We analyzed all theoretically possible codon substitutions that could have occurred by single point mutations in 520 cancer census genes and calculated their mutability values based on their genomic context. We found that only about one percent of all theoretically possible codon substitutions were observed in the surveyed 12,013 tumor samples derived from the COSMIC v85 cohort (Table S1). Using the pan-cancer model, across all analyzed possible codon substitutions produced by single point mutation, mutability ranged from 1.61 × 10^−7^ to 1.80 × 10^−5^ (mean = 1.34 × 10^−6^). Lower and upper boundaries for mutability are dependent on the cancer model selection, and cancer models with higher mutational burdens like melanomas (1.92 × 10^−7^ to 1.35 × 10^−4^, mean = 7.00 × 10^−6^) have higher mutability values compared to cancers such as prostate adenocarcinoma (5.12 × 10^−8^ to 7.31 × 10^−6^, mean = 3.95 × 10^−7^).

We found that across codon substitutions which were not observed in the COSMIC v85 cohort, the mean mutability (1.29 × 10^−6^) was found to be three-fold lower compared to the mutability of observed codon substitutions (3.88 × 10^−6^) using pan-cancer background model, Mann-Whitney-Wilcoxon test *p* < 0.01 (Figure 1A). This finding also holds true for different cancer-specific models (the list of cancer-specific mutational profiles can be found in https://www.ncbi.nlm.nih.gov/research/mutagene/signatures#mutational_profiles). The same result is confirmed for per-nucleotide mutability (1.04 × 10^−6^ versus 3.36 × 10^−6^, Mann-Whitney-Wilcoxon test *p* < 0.01). In addition, we validated our result on a set of observed mutations from 9,228 patients who had undergone prospective sequencing of MSK-IMPACT gene panel. Looking at mutations in the genes which were sequenced in all patients in the MSK-IMPACT cohort, the same pattern remains that observed codon substitutions had a higher mutability (3.41 × 10^−6^), compared to those which were theoretically possible, but did not occur in cancer patients (1.30 × 10^−6^, Mann-Whitney-Wilcoxon test, *p* < 0.01) (Figure 1B).

**Figure 1.**
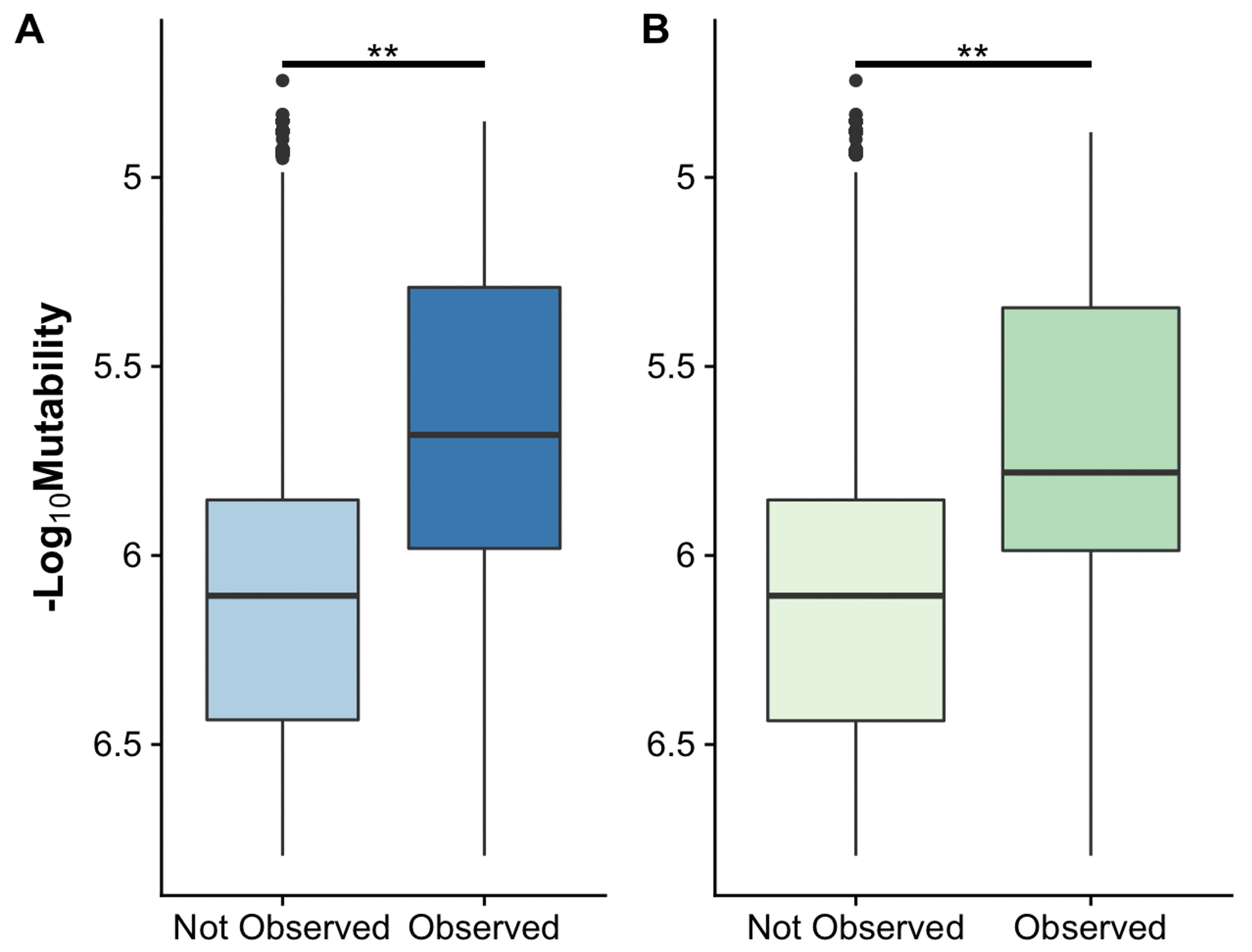
Mutability of all theoretically possible codon substitutions (“not observed”) and all substitutions that were observed in: **(A)** COSMIC v85 pan-cancer cohort; **(B)** MSK-IMPACT cohort. Asterisks show the differences on Mann-Whitney-Wilcoxon test significant at p < 0.01. Mutability values have been converted to negative log_10_ scale as pan-cancer codon mutability ranges several orders of magnitude.

Figure S1 shows cumulative and probability density distributions of nucleotide mutability values for all observed mutations in patients, for theoretically possible mutations in all cancer census genes and for two genes in particular, CASP8 and TP53. While there are many theoretically possible mutations with low mutability values, the observed cancer spectrum is dominated by mutations with high mutability. A similar pattern is seen for cancer-specific cases (Figure 2).

**Figure 2.**
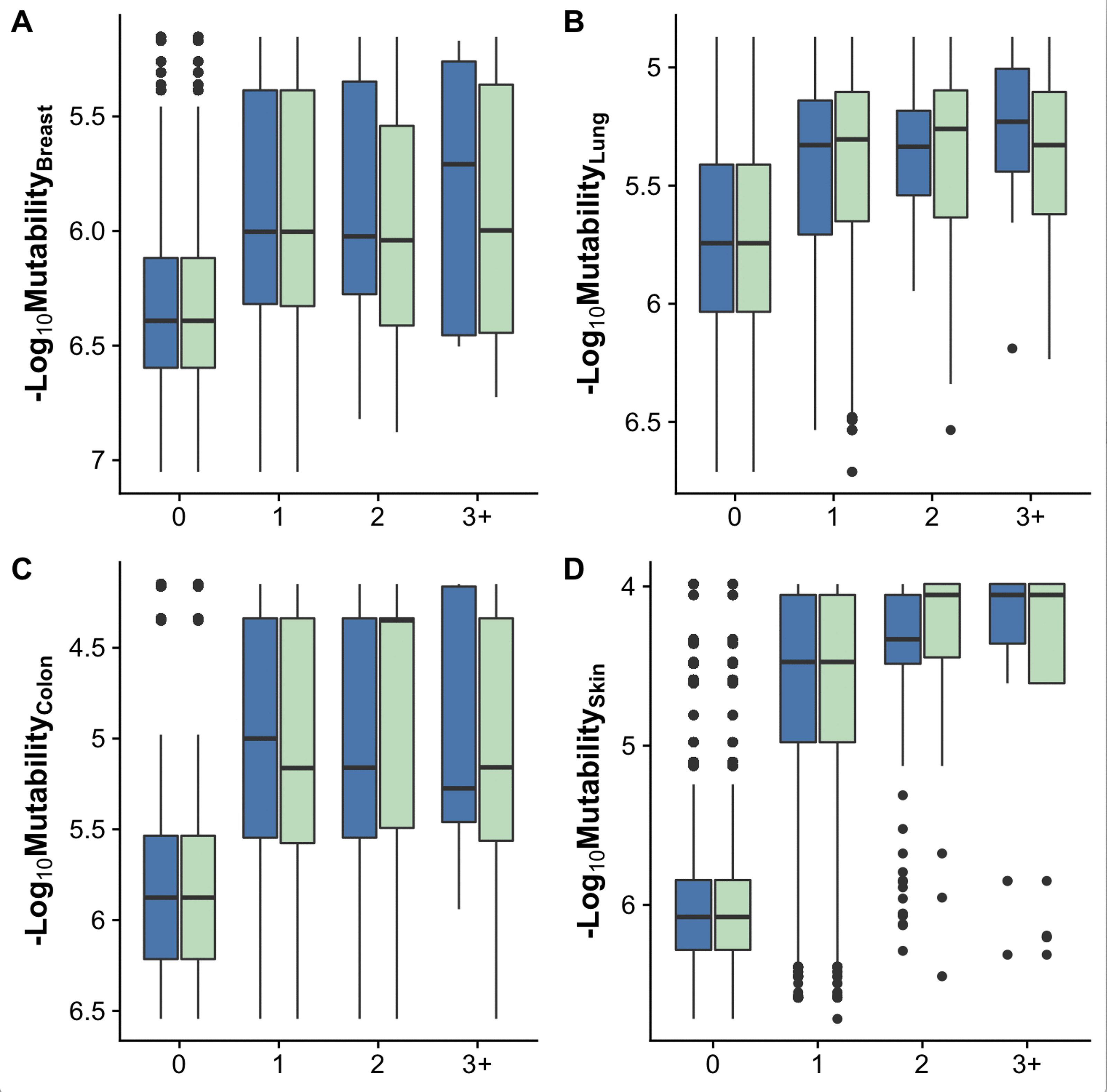
Relationship between cancer-specific nucleotide mutability and observed reoccurrence frequency of all mutations from two cohorts. Counts are binned and refer to how many times a particular mutation was observed in the given cancer type. ‘0’, ‘1’, ‘2’ and ‘3+’ refer to mutations that were not observed (including all possible point mutations), observed once, twice, or in three or more cancer samples. Blue boxes show mutations with the observed frequency calculated in the COSMIC v85 cohort and green boxes refer to MSK-IMPACT cohort. **(A)** breast cancer (n_COSMIC_ = 1,667, n_MSK_ =783 samples), **(B)** Lung adenocarcinoma (n_COSMIC_ = 301, n_MSK_ = 1,203), **(C)** Colon adenocarcinoma (n_COSMIC_ = 369, n_MSK_ = 688) and **(D)** Skin malignant melanoma (n_COSMIC_ = 376, n_MSK_ =182).

### Silent mutations have the highest mutabilities

Figures 3A,B show the distributions of codon mutability values for all possible missense, nonsense, and silent mutations accessible by single nucleotide base substitutions in the protein-coding sequences of 520 cancer census genes calculated with the pan-cancer background model. Codon mutability spans two orders of magnitude, and silent mutations have significantly higher average mutability values (mean = 5.68 × 10^−6^) than nonsense (mean = 3.44 × 10^−6^) or missense mutations (mean = 3.29 × 10^−6^) (Kruskal-Wallis test *p* < 0.01 and Dunn’s post hoc test *p* < 0.01 for all comparisons). These differences in *codon mutabilities* could be a reflection of the degeneracy of genetic code, where multiple silent nucleotide substitutions in the same codon may increase its mutability. However, degeneracy of genetic code does not affect the calculation of *nucleotide mutability*. While the differences between types of mutations are less pronounced for nucleotide mutability (Figure 3C), silent mutations are still characterized by the highest nucleotide mutability values (mean = 3.91 × 10^−6^ for silent, 3.10 × 10^−6^ for nonsense and 3.17 × 10^−6^ for missense mutations, Kruskal-Wallis test *p* < 0.01 and Dunn’s post hoc test *p* < 0.01 for all comparisons).

**Figure 3.**
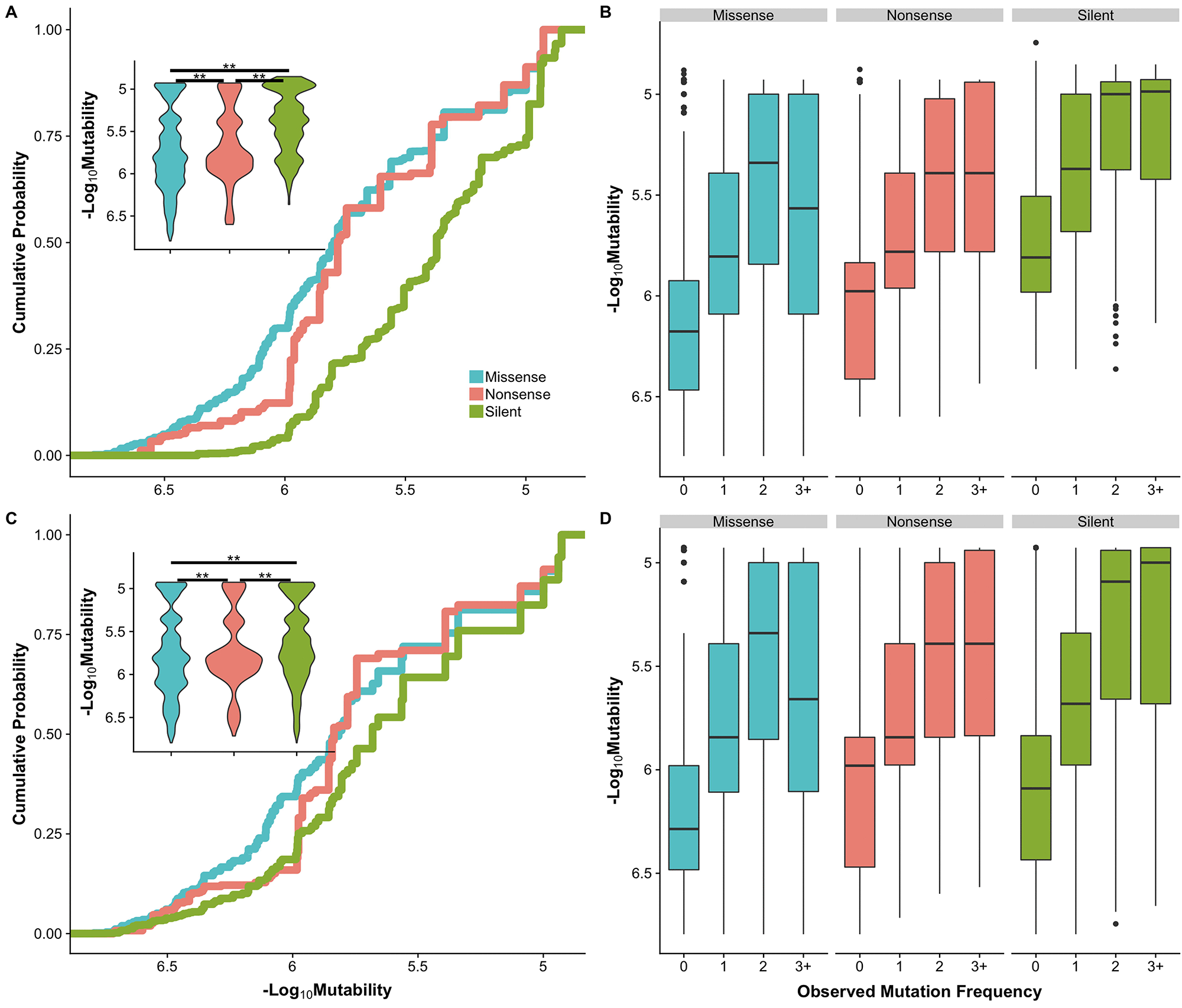
Mutability distributions by mutation type and mutation frequency. **(A)** Cumulative distribution of *codon mutability* of silent (green), nonsense (red) and missense (blue) mutations. **(C)** Cumulative distribution of *nucleotide mutability* for silent, nonsense and missense mutations. Inset shows the probability density distributions of mutability by mutation type. Significance was determined by Dunn’s test; difference with *p* < 0.01 is marked with a double asterisk. **(B)** and **(D)** are codon and nucleotide mutability respectively binned by frequency in the COSMIC v85 pan-cancer cohort. ‘0’, ‘1’, ‘2’ and ‘3+’ refer to mutations that were not observed (including all possible point mutations), observed once, twice, or in three or more cancer samples. See Table S1 for the number of mutations in each category.

### Background mutability significantly contributes to shaping the observed mutational spectrum

Under the null model of all mutations arising as a result of background mutational processes, somatic mutations should accumulate with respect to their mutation rate and one would expect a positive correlation between mutability and observed mutational frequency of individual mutations. Indeed, as Figures 3B,D show, this is the case for silent and nonsense mutations. To further investigate this relationship, in the pan-cancer COSMIC v85 cohort we calculated both Spearman’s rank, a non-parametric test taking into account that mutability is not normally distributed, and Pearson linear correlation coefficients between codon mutability and frequencies of mutations across all 520 cancer census genes. We also explored this association for each gene with at least ten unique mutations of each type: silent, nonsense, and missense (Figure 4).

**Figure 4.**
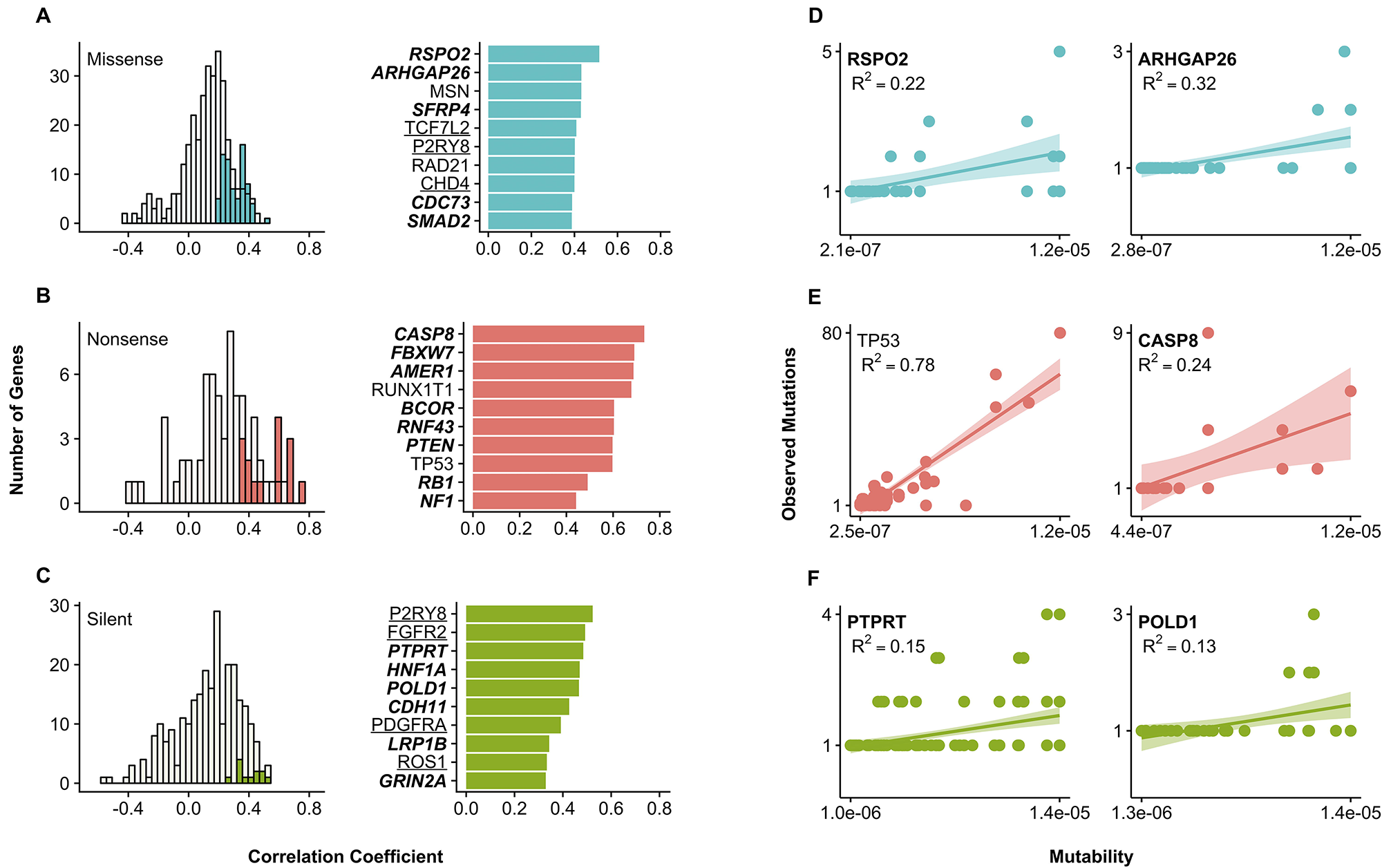
Relationship between codon mutability and frequency of mutations. Histograms show the Spearman rank correlation coefficients between the reoccurrence frequency and mutability across cancer genes with at least 10 observed mutations of each type: **(A)** missense (blue), **(B)** nonsense (red) and **(C)** silent (green). Filled bars in the left column denote genes with significant correlation at *p* < 0.01. Bar graphs show Spearman correlation coefficient for genes with significant correlation at *p* < 0.01. Genes with bold font are tumor suppressors (TSG), underlined genes are oncogenes, and genes in plain font were either categorized as both TSG and oncogene or fusion genes. **(D-F)** Scatterplots with regression lines and confidence intervals show the linear relationship between mutability and reoccurrence frequency of each type of mutation for several representative genes. Adjusted R^2^ are shown to convey goodness of fit. Mutation reoccurrence frequencies were taken from the pan-cancer COSMIC v85 cohort.

Overall, we found 84 and 137 genes with significant (*p* < 0.01) positive Spearman and Pearson correlations, respectively, for at least one mutation type (Table S2). Among the genes with significant correlations, 41 belong to tumor suppressor genes, 28 are oncogenes, and 15 genes are classified as either fusion genes or both oncogene and tumor suppressor. For some genes, including TP53 (first column, Figure 4E) and tumor suppressor CASP8 (second column, Figure 4E), a strong linear relationship between mutability and recurrence frequency of observed mutations (*R*^2^ > 0.5) was observed. Breaking up all codon changes into silent, nonsense and missense reveals the highest correlations for silent (*p* = 0.15, *r* = 0.1, *p* < 0.01) and nonsense (*p* = 0.20, *r* = 0.15, *p* < 0.01) mutations (Figure S2).

### Relationship between mutability and observed frequency is different for tumor suppressor and oncogenes

The effects of mutations on protein function, with respect to their cancer transforming ability, can drastically differ in tumor suppressor genes (TSG) and oncogenes, therefore we performed our analysis separately for these two categories (Figure 5). In general, mutations in TSG can cause cancer through the inactivation of their products, whereas mutations in oncogenes may result in protein activation. We used COSMIC gene classification separating genes into tumor suppressors and oncogenes. Genes which were annotated as both TSG and oncogenes were excluded from this analysis. Gene ontology (GO) analysis found that top GO annotations in TSG for cellular compartments were “nucleus”, “chromosome”, and “nuclear part” and for molecular functions were “protein”, “DNA”, and “enzyme binding”. For the oncogenes, the top GO annotations for cellular components were “nucleoplasm”, “nucleus”, and “nuclear lumen” and for molecular function “heterocyclic compound binding”, “organic cyclic compound biding” and “sequence-specific DNA binding”. A full list of genes and the associated GO terms is available in Supplemental Table S3. In addition, we used COSMIC classification into genes with dominant or recessive mutations, but overall results were similar to the ones produced using classification into TSG and oncogenes (Figure S3).

**Figure 5.**
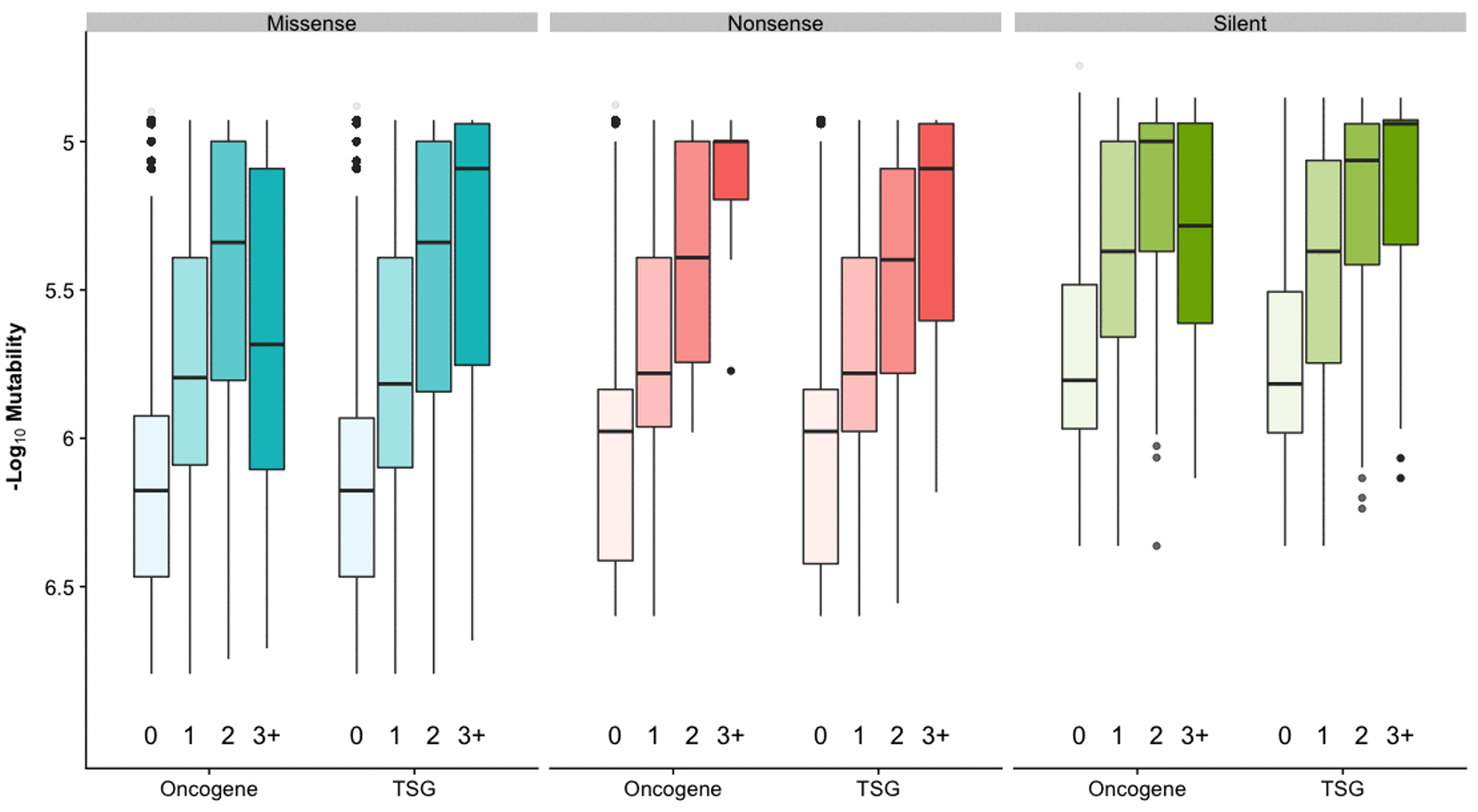
Relationship between codon mutability and reoccurrence frequency of mutations for different mutation types and gene functions. Genes grouped into oncogene and tumor suppressor (TSG) by their role in cancer. Mutations were binned by their reoccurrence frequency in COSMIC v85 cohort. Boxplots show codon mutability calculated with pan-cancer model. See Table S1 for counts.

We observed a weak but statistically significant correlation between codon mutability and recurrence frequency in TSG (*p* = 0.17, *r* = 0.13, *p* < 0.01) while oncogenes showed a weaker Spearman correlation and no significant Pearson correlation (*p* = 0.13, *p* < 0.01; *r* = 0, *p* = 0.61) (Figure S2B,C). This correlation mostly arises from neutral mutations as shown in the following section. An inverse U-shaped trend was detected for missense and silent mutations in oncogenes: highly recurrent mutations (observed in three and more samples) were characterized by low average mutability values (Figure 5). In the latter case, selection may be a more important factor compared to background mutation rate explaining reoccurrence of these mutations. Functionally conserved sites overall were found to be more frequently mutated in oncogenes [29], and our analysis did not find a straightforward association between mutability and evolutionary conservation.

### Neutral mutations have higher mutability values than non-neutral

We complied a *combined* dataset of experimentally annotated missense mutations in cancer genes from several sources. Mutations were categorized as ‘non-neutral’ or ‘neutral’ based on their experimental effect on protein function, transforming effects, and other characteristics (see Methods and Table S4). For all mutations in *combined* dataset, whether they were observed in MSK-IMPACT or the COSMIC v85 cohorts, the codon mutability values of neutral mutations were significantly higher (mean = 2.71 × 10^−6^) (Mann-Whitney-Wilcoxon test, *p* < 0.01) than for non-neutral mutations (mean = 1.74 × 10^−6^) (Figure 6A). Binning the mutations by their reoccurrence frequency also showed differences between ‘neutral’ and ‘non-neutral’, with the frequency of neutral mutation depending on their mutability. For neutral mutations, mutations that were observed in three or more samples had higher background mutability (mean_MSK_ = 6.39 × 10^−6^, mean_COSMIC_ = 6.22 × 10^−6^) compared to mutations which were not observed (mean_MSK_ = 2.46 × 10^−6^, mean_COSMIC_ = 2.54 × 10^−6^). In contrast, the background mutability of non-neutral mutations did not vary with the reoccurrence frequency (Figure 6B), suggesting that background mutability was much less important in driving reoccurrence of non-neutral mutations.

**Figure 6.**
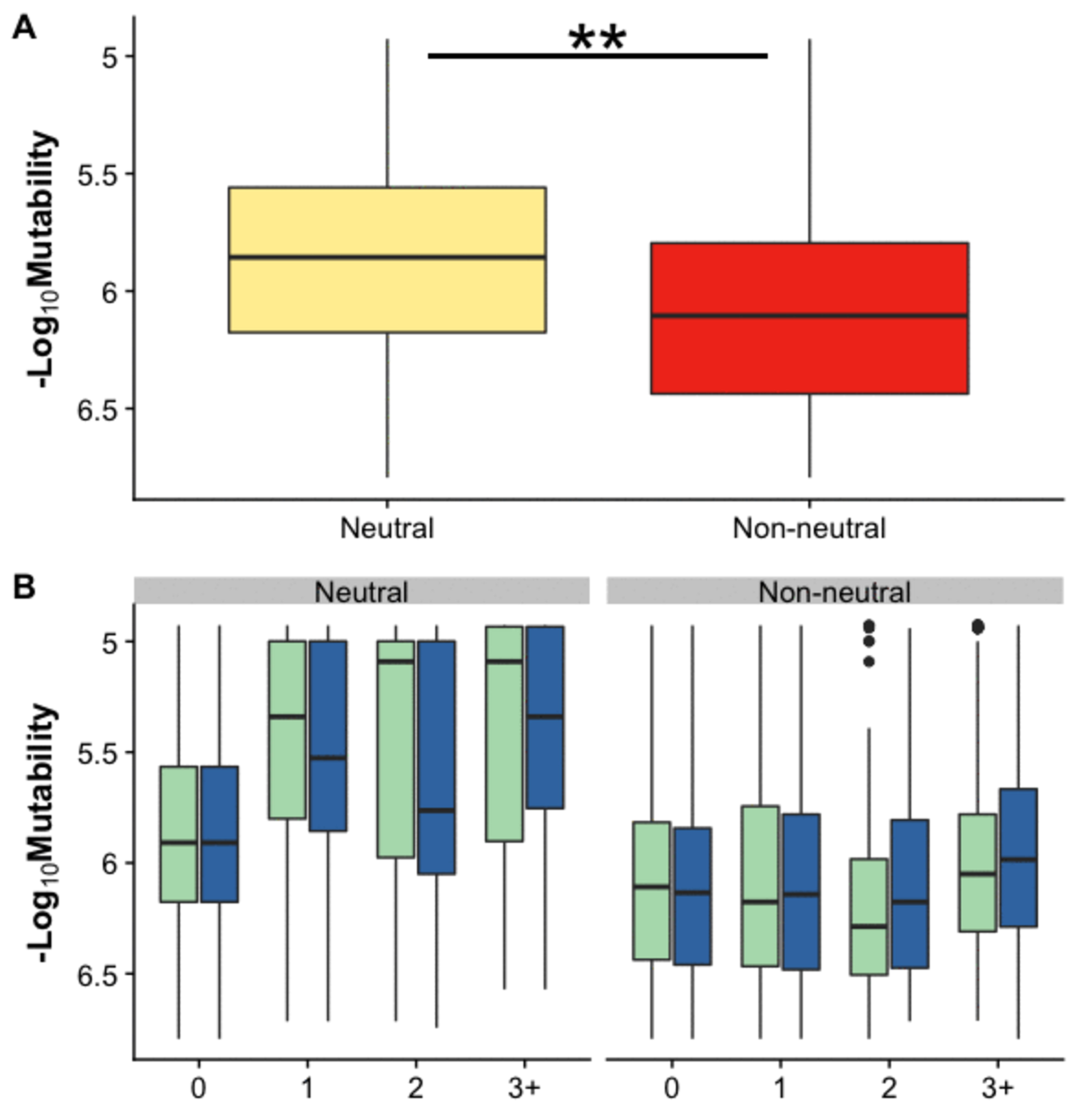
Codon mutability of missense mutations grouped by their experimental effects. **(A)** Mutations from the *combined* dataset were categorized as neutral and non-neutral. Significant differences with *p* < 0.01 are marked with a double asterisk. Mutability was calculated with pan-cancer background model **(B)** Mutations binned by their reoccurrence frequency in both MSK-IMPACT (green) and COSMIC v85 (blue) cohorts. In both cohorts, reoccurrence frequency of neutral mutations depends on mutability, whereas for non-neutral mutations, reoccurrence frequency does not scale with background mutability.

### Accounting for context-dependent mutability in ranking of mutations

In the previous sections we explored the contribution of background mutational processes in understanding the observed mutational patterns in cancer. With our finding that background mutability differs between neutral mutations and non-neutral mutations, we explored if background mutability could be used to facilitate the detection of cancer driver mutations or provide a reasonable ranking in terms of their potential carcinogenic effects. We tested different ways to calculate codon mutability and if it could help to differentiate between experimentally annotated neutral, or putatively passenger mutations, and non-neutral driver mutations. We found that a simple and intuitive measure, B-score, calculated without using gene weights (see next section) performed the best on the *combined* experimental test set. A similar measure was used previously to identify mutational hot spots [21, 30]. Hotspots are defined for sites, whereas our approach assesses specific mutations, and different mutations from the same hotspot can be drivers or passengers. For instance, TP53 Tyr236 site is annotated as a hotspot in [21, 30], however p.Tyr236Phe mutation in this site is experimentally characterized as neutral in the IARC database.

We compared the performance of B-score to six state-of-the-art computational methods which distinguish driver from passenger mutations in cancer: CHASM [31], CHASMplus [32]VEST[33], REVEL[34], CanDrAplus[35]and FatHMM[36]. Table 1 shows the performance of the various computational predictors at classifying mutations from the *combined* dataset observed in two sets of cancer cohorts. To compare across methods, which use different thresholds for calling neutral versus non-neutral mutations, we calculated the Matthew’s correlation coefficient (MCC) across a range of thresholds for each method and reported the maximal MCC value. Based on the MCC, the best classifiers are CHASMplus, B-score and CanDrAplus (MCC = 0.64, 0.61, and 0.58 respectively) (Table 1). Surprisingly, mutation reoccurrence frequency alone performs very well, with MCC of 0.49 in the COSMIC v85 cohort and 0.51 in the MSK-Impact cohort. B-Score is able to provide a correction to reoccurrence frequency using codon mutability and yields a much better performance than frequency alone. Intriguingly, inverse mutability alone performs better than random, emphasizing the fundamental quality of non-neutral mutations in cancer: mutability of driver mutations is lower than the mutability of passengers (Figure 6).

**Table 1.**
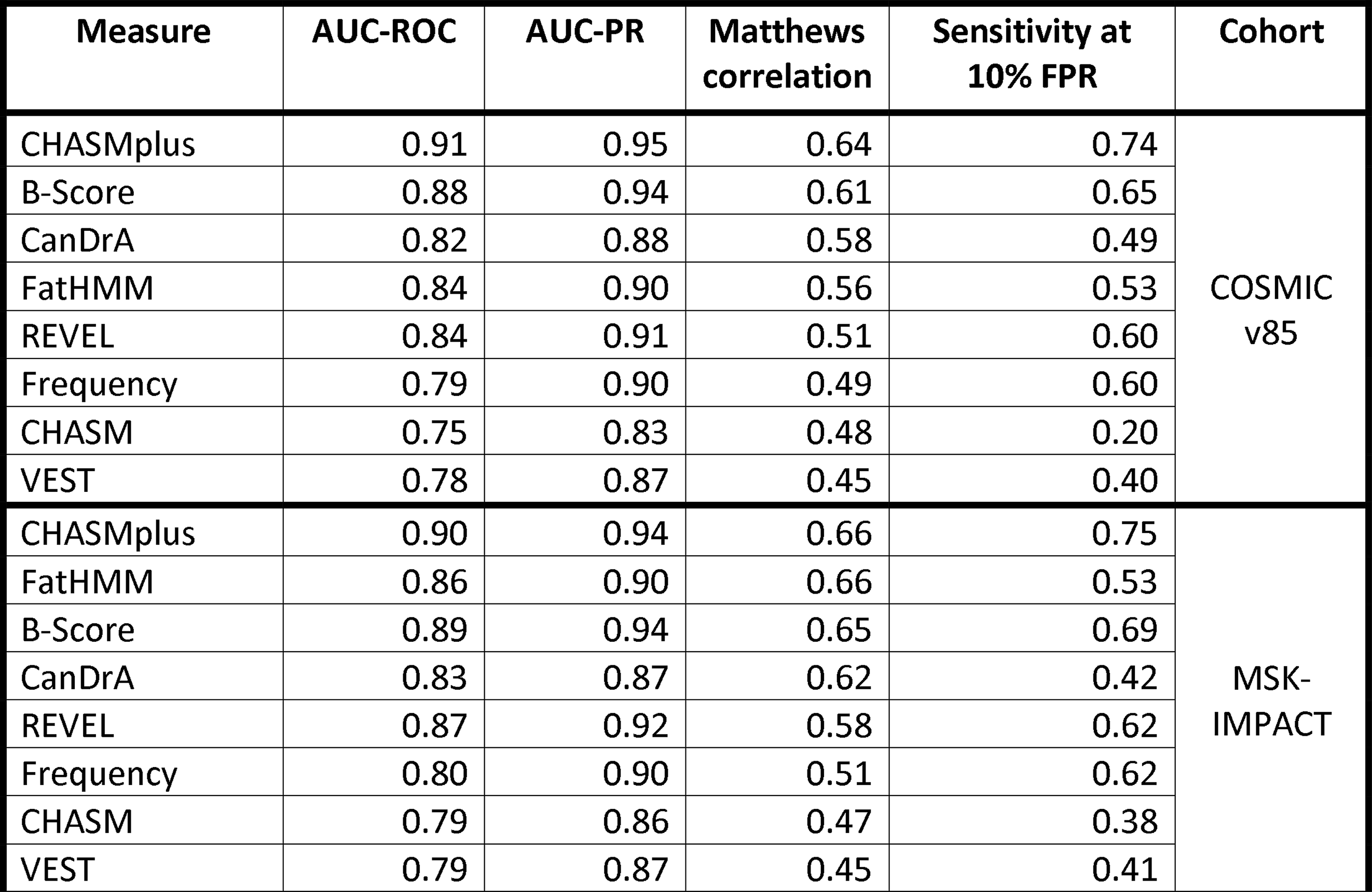
Comparison of different methods to distinguish neutral from non-neutral mutations. from *combined* experimental dataset. Mutations were observed in corresponding cancer cohorts. See Table S6 and S7 for results on rare and all mutations. Maximum Matthew’s correlation is reported for each predictor. B-Score for each cohort is calculated with the respective cohort size: COSMIC v85 cohort 12,013; MSK-Impact 9,228. For CHASM the background model yielding best performance was chosen.

We also explored the performance of methods in classifying mutations that were not observed or observed only once in the COSMIC v85 cohort or MSK-Impact cohort (Table S6). For mutations which were not observed in the COSMIC v85 cohort B-Score classification performance is low but better than random (AUC=0.65). On mutations which were observed in only one cancer sample in the cohort (207 passenger and 157 driver mutations), B-Score still performed better than VEST and CHASM (MCC = 0.46, 0.42, and 0.36 respectively). On the combined set which includes all experimentally verified mutations, whether they were observed or not observed in cancer patients, B-score ranks fourth after CHASMplus, REVEL and FatHMM (Table S7).

B-score also allows to break ties for mutations observed in the same number of patients. For example in the TP53 gene, mutations p.Glu11Lys and p.Cys135Gly have been observed in two patients each in the COSMIC v85 cohort. However, p.Glu11Lys (mutability of 1.18 × 10^−5^) is predicted a passenger mutation and p.Cys135Gly (mutability of 2.20 × 10^−7^) is predicted as a driver mutation which is consistent with the annotations from the experimental *combined* dataset.

### Variability of mutation rates across genes

Even though our probabilistic model indirectly incorporates different factors affecting mutation rate, we checked explicitly if large-scale factors, allowing mutations of the same type to have different mutational probabilities in different genes, affected retrieval performance on the *combined* test set. Several methods have been developed to estimate gene weights (see Methods), which consider the overall number of mutations or the number of silent mutations affecting a gene. Additionally, we estimated the gene weights based on the number of SNPs in the vicinity of a gene. We also examined the effects of several large-scale confounding factors such as gene expression levels, replication timing, and chromatin accessibility (provided in the gene covariates in MutSigCV [37]) on gene weights. We used gene weights to adjust mutability values and explored whether any of the gene weight models were helpful in distinguishing between experimentally determined neutral and non-neutral mutations. We found that “no-outlier”-based weight (*r* = 0.66, *p* = 0.004) and “silent mutation”-based weight (*r* = 0.65, *p* = 0.004) significantly correlated with the gene expression levels. No other correlations of gene weights with confounding factors were found. Overall, using gene weight as an adjustment for varying background mutational rates across genes did not improve classification performance of mutations in the experimental benchmark. Only a SNP-based weight affected the AUC-ROC, but the gain was very minimal, and no gene weight affected the MCC (Table S8). One of the reasons could be the limited number of genes used in the experimental benchmark set (58 genes).

### Ranking of cancer point mutations in MutaGene

MutaGene webserver provides a collection of cancer-specific context-dependent mutational profiles [38] It allows to calculate nucleotide and codon mutability and B-Score for missense, nonsense and silent mutations for any given protein coding DNA sequence and background mutagenesis model using the “Analyze gene” option. Following the analysis presented in this study, we added options to provide a ranking of mutations observed in cancer samples based on the B-Score or the multiple-testing adjusted q-values. Using the *combined* dataset as a performance benchmark (Table 1, Table S7), we calibrated two thresholds: the first corresponds to the maximum of MCC, and the second corresponds to 10% FPR. Mutations with the B-Score below the first threshold are predicted to be “cancer drivers”, whereas mutations with scores in between two thresholds are predicted to be “potential drivers”. All mutations with scores above the second threshold are predicted as “passengers”. Importantly, calculations are not limited to pan-cancer and can be performed using a mutational profile for any particular cancer type, the latter would result in a cancer-specific ranking of mutations and could be useful for identification of driver mutations in a particular type of cancer. An example of prediction of driver mutations status for EGFR is shown in Figure 7. MutaGene Python package allows to rank mutations in a given sample or cohort in a batch mode using pre-calculated or user-provided mutational profiles and available at https://www.ncbi.nlm.nih.gov/research/mutagene/gene.

**Figure 7.**
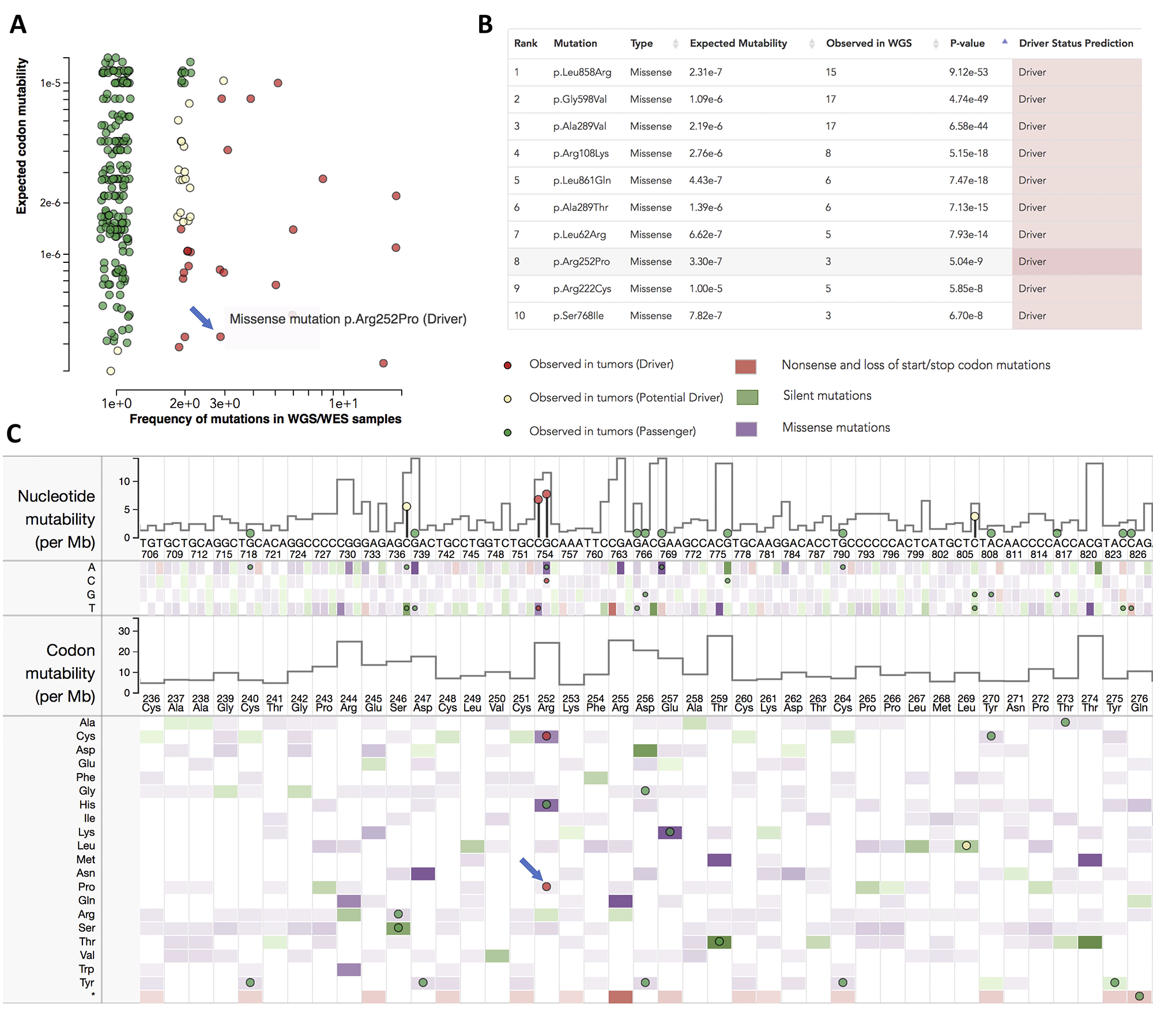
Ranking of mutations and prediction of driver mutations based on B-score. Snapshots from the MutaGene server show the results of analysis of EGFR gene with a Pan-cancer model. **(A)** Scatterplot with expected mutability versus observed mutational frequencies. **(B)** Top list of mutations ranked by their B-Scores. **(C)** EGFR nucleotide and translated protein sequence shows per-nucleotide site mutability per codon mutability as well as mutabilities of nucleotide and codon substitutions (heatmaps). Mutations observed in tumors from ICGC repository are shown as circles colored by their prediction status: Driver, Potential driver, and Passenger. Missense mutation p.Arg252Pro is shown with a blue arrow.

## Discussion

To understand what processes drive point mutation accumulation in cancer, we used DNA context-dependent probabilistic models to estimate the baseline mutability for nucleotide mutation or codon substitution in specific genomic sites. Passenger mutations, constituting the majority of all observed mutations, may have largely neutral functional impacts and are unlikely to be under selection pressure. For passenger mutations one would expect that mutations with lower DNA mutability would have lower observed mutational frequency and vice versa. In a recent study the fraction of sites harboring SNPs in the human genome was indeed found to correlate very well with the mutability although the later was estimated differently from our study [39]. We detected a significant positive correlation between background mutability, which is proportional to per-site mutation rate, and observed reoccurrence frequencies of mutations in cancer patients. In accordance with this trend, we also found that mutations that were not observed in cancer cohorts were marked by a lower background mutability. For some genes, such as TP53 or CASP8, mutations and their frequencies can be predicted from their mutability values. Outliers of this association trend, mutations which reoccur at high frequencies but have low mutabilities might be important for inferring mutations under positive selection, as illustrated especially for missense mutations in oncogenes.

In this respect, reoccurring synonymous mutations with low mutability may represent interesting cases for further investigation of potential synonymous drivers. Mutability of synonymous mutations was found to be the highest among other types of mutations. Observed mutational frequency of synonymous mutations scales with their mutability, therefore it is important to correct for mutability while ranking these mutations with respect to their driver status. Overall, B-score predicted 102 synonymous driver mutations in 64 out of 520 cancer-associated genes. It has been previously shown that some synonymous mutations might be under positive selection and can affect the speed and accuracy of transcription and translation, protein folding rate, and splicing [40]. Some recurrent highly mutable synonymous mutations might not represent relevant candidates of drivers, whereas some rare mutations with relatively low mutability are predicted to be drivers by our approach (e.g. KDR gene p.Leu355=, NTRK1 gene p.Asn270=).

In this paper we developed and tested a probabilistic model, implemented as B-Score, to adjust the reoccurrence frequency of a mutation (a measure commonly used in clinical research to identify genes and mutations under selection) by its expected background mutability. B-Score is able to provide a correction to reoccurrence frequency using mutability and improves the classification of cancer driver and passenger mutations by up to 20% compared to reoccurrence frequency alone. The advantages of B-score are that: (i) it is intuitive and interpretable, (ii) does not rely on many parameters, and (iii) does not involve explicit training on driver and passenger mutation sets. One of the disadvantages is that it requires the knowledge of a total number of patients tested. We found that B-Score performed comparably or better to many of state-of-the-art methods even for rare mutations observed in two large cancer cohorts. However, it underperformed for those mutations from combined experimental set that were not observed in cancer patients. These latter mutations might either constitute functionally disruptive mutations not directly connected with the carcinogenesis or might represent rare cancer mutations not yet detected in large cancer cohorts.

A lot of efforts have been focused so far on developing a comprehensive set of cancer driver mutations verified at the levels of functional assays or animal models [26, 41, 42]. However, existing sets often contain predictions and very few neutral cancer passenger mutations. The vast majority of computational prediction methods rely on machine learning algorithms trained on mutations from a few genes and/or on recurrent mutations as estimates of driver events or use germline SNPs or silent mutations as the presumed “neutral” set. In many cases, the performance is evaluated on similarly generated synthetic benchmarks. As a result, methods can be trained on incorrectly labeled data and even if trained on correct data, can exhibit a well-known overfitting effect.

While mutational processes vary widely among cancer types, and different driver mutations have been shown to be preferentially associated with specific mutational processes [39, 40], there remains a lack of cancer-specific driver/passenger datasets. In our combined dataset, the effects of mutations were determined using experimental assays, which were not linked to any particular cancer type, therefore a pan-cancer model was used for calculation of B-score and other methods tested. However, we provide the ability to apply cancer-specific B-Score ranking of mutations using the models available via the MutaGene package and website (see Methods). Additionally, for some cancer types, the background mutational processes may differ greatly between subsets of cancer patients. For these highly heterogenous cancer types rather than using cancer type specific, it may be more appropriate to use background mutational profiles/models specific for a given cohort.

In this study, we restricted our test dataset to only missense mutations that have been experimentally assessed, with several thousands of driver and passenger mutations from 58 genes. Intriguingly, we found that experimentally annotated driver mutations had a lower background mutability than neutral mutations, suggesting possible action of context-dependent codon bias towards less mutable codons at critical sites for these genes, although more studies would have to be conducted to further investigate this observation. This important difference in mutability between drivers and passengers may explain the outstanding performance of the simple measure B-score which enables an understanding of the differential roles that background mutation rate and selection play in shaping the cancer mutational spectrum.

## Methods

### Defining driver and passenger mutations using datasets of experimental assays

We assembled a *combined dataset* that included mutations from the five datasets described below. First we obtained missense mutations for TP53 gene with experimentally determined functional transactivation activities from IARC P53 database where they were classified as functional, partially-functional, and non-functional[43].

The second dataset contained experimental evidence collected from the literature[44]. The experimental evidence of impact of mutations included changes in enzymatic activity, response to ligand binding, impacts on downstream pathways, an ability to transform human or murine cells, tumor induction *in vivo*, or changes in the rates of progression-free or overall survival in pre-clinical models. Mutations were considered “damaging” if there was literature evidence to support their impact on at least one of the above-mentioned categories. Mutations with no significant impacts on the wild-type protein function were classified as “neutral”. Mutations with no reliable functional evidence were regarded as “uncertain” and were not used in this study.

The third dataset included experimentally verified BRCA1 mutations and was originally collected by using deep mutational scanning to measure the effects of missense mutations on the ability of BRCA1 to participate in homology-directed repair. In this dataset missense mutations were categorized as either “neutral” or “damaging” [45, 46]. Noteworthy, BRCA1 set contained inherited germline as well as somatic mutations.

The fourth dataset explored over 81,000 tumors to identify drivers of hypermutation in DNA polymerase epsilon and polymerase delta genes (POLE/POLD1). “Drivers of hypermutation” were variants which occurred in a minimum of two hypermutant tumors, which were never found in lowly mutated tumors, and did not co-occur with an existing known driver mutation in the same tumor. Other variants in these genes were considered “passengers” with respect to hypermutation[25].

The fifth dataset consisted of missense mutations annotated based on their effects on cell-viability in Ba/FC and MCF10A models[47]. “Activating mutations” were mutations where the cell viability was higher than the wild-type gene, and “neutral mutations” were those mutations where cell-viability was similar to the wild-type. Ng *et al.* used these consensus functional annotations to compare the performance of 21 different computational tools in classifying between activating and neutral mutations using ROC analysis, with activating mutations acting as the positive set and neutral as the negative set. The authors found that the tools yielding best performance were CanDrAplus and CHASM. We included 743 mutations (488 neutral and 255 activating) in 50 genes accessible through single nucleotide substitutions out of the 816 activating and neutral mutations that Ng et al tested [47].

Finally, we removed redundant and conflicting entries when mutations annotated as non-functional or neutral in one dataset were also annotated as damaging or benign in another. As a result, all mutations in the *combined* data set were categorized as “non-neutral” (affecting function, binding or transforming) and “neutral” (other mutations). We treated “functional” and “partially -functional” mutations in IARC TP53 dataset as “neutral”, and “non-functional” as “non-neutral”. Overall, the *combined dataset* contains 5,276 mutations (4,137 neutral and 1,139 non-neutral) from 58 genes (Table S4) and is available on MutaGene website at https://www.ncbi.nlm.nih.gov/research/mutagene/benchmark.

### Datasets of mutations observed in cancer patients

The Catalogue of Somatic Mutations in Cancer (COSMIC) database stores data on somatic cancer mutations and integrates the experimental data from the full-genome (WGS) and whole-exome (WES) sequencing studies [48]. Cancer census genes (520 genes) were defined according to COSMIC release v84. For each of these genes, we explored all theoretically possible nucleotide mutations along the DNA sequence of the principal transcripts. This resulted in 4,129,461 possible nucleotide substitutions, and 3,293,538 codon substitutions.

For analyses comparing oncogenes and tumor suppressor genes (TSG), genes classified as only fusion genes or those with both oncogenic and TSG activities were not used. This resulted in 205 oncogenes and 167 TSG (Table S3). For gene ontology (GO) enrichment analysis we used the R package “GOfuncR”. For enrichment analysis, the genes annotated as either TSG or oncogenes were compared to all other genes in the “Homo.sapiens” gene annotation package in R. 98% of all COSMIC v85 samples contained less than 1000 mutations so were not hypermutated. COSMIC v85 samples which came from cell-lines, xenografts, or organoid cultures were excluded. Only mutations with somatic status of “Confirmed somatic variant” were included and mutations which were flagged as SNPs were excluded.

For each cancer patient, a single sample from a single tumor was used. Additionally, it is possible that the same patient may be assigned different unique identifiers in different papers, and duplicate tumor samples are sometimes erroneously added to COSMIC database during manual curation. These samples may affect the recurrence counts of mutations. We applied clustering method in order to detect and remove any redundant tumor samples. Each sample was represented as a binary vector with 1 if a sample had a mutation in a particular genomic location and 0 otherwise. The binary vectors were compared with Jaccard distance metric, 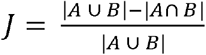, where identical samples have *J* = 0, followed by agglomerative clustering with complete linkage. Non-singleton clusters with pairwise distance cutoff of *J* ≥ 0.3 were extracted and only one representative of each cluster was used, whereas other samples were discarded. Because of these relatively stringent criteria for inclusion, it is likely that some small number of non-duplicate samples were discarded in this process.

MSK-IMPACT cohort was obtained from cBioPortal [49]. We ensured that no mutations were counted multiple times for each patient; if there were multiple tumor samples per patient, primary and metastatic, the primary tumor was kept, and the metastatic discarded. Only those tumors which were sequenced against a matched normal sample were kept to ensure validity of somatic mutations.

In the 520 genes we explored, we investigated if these genes were expressed in cancer cell lines from multiple tissue types using RNAseq data from the January 2019 release of the Cancer Cell Line Encyclopedia [50]. Using RNAseq data of 1,019 unique cancer cell lines from 26 different tissue types and a cutoff for expression at 0.5 RPKM (Reads Per Kilobase of transcript, per Million mapped reads), we found that 512 genes were expressed in at least one tissue.

### Calculation of context-dependent DNA background mutability

Context-dependent mutational profiles were constructed previously from the pools of mutations from different cancer samples by counting mutations observed in a specific trinucleotide context [38]. Altogether, there are 64 different types of trinucleotides and three types of mutations *x*≫*y* (for example C≫A, C≫T, C≫G and so on) in the central position of each trinucleotide which results in 192 trinucleotide context-dependent mutation types. In a mutated double-stranded DNA both complementary purine-pyrimidine nucleotides are substituted, and therefore we considered only substitutions in pyrimidine (C or T) bases, resulting in 96 possible context-dependent mutation types *m* = *a*[*x* ≫ *y*]*b*, where *a*, *b*, *x*, *y* ∈ {*A*, *T*, *C*, *G*}, *x* ≠ *y*. Thus, mutational profile can be expressed as a vector of a number of mutations of certain type (*f*_l_, …, *f*_96_) or a number of mutations of certain type per sample (*r*_l_,…, *r*_96_). Profiles were constructed under the assumption that vast majority of cancer context-dependent mutations have neutral effects, while only a negligible number of these mutations in specific sites are under selection. To assure this, we removed recurrent mutations (observed twice or more times in the same site) as these mutations might be under selection in cancer. In the current study we used pan-cancer and cancer-specific mutational profiles for breast, lung adenocarcinoma, colon adenocarcinoma, and skin melanoma derived from MutaGene [38].

We calculated mutability that described baseline DNA mutagenesis per nucleotide or per codon. ***Mutability*** was defined as a probability to obtain a context-dependent nucleotide mutation purely from the baseline mutagenic processes operating in a given group of samples. Mutability is proportional to the expected mutation rate of a certain type of context-dependent mutation regardless of the genomic site it occurs. For exome mutations, given the number of different trinucleotides of type *t* in a diploid human exome, *n_t_*, the ***nucleotide mutability*** is calculated as:

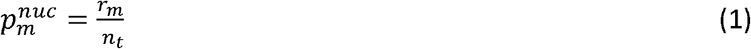

In protein-coding sequences it is practical to calculate mutation probability for a codon in its local pentanucleotide context, given trinucleotide contexts of each nucleotide in the codon. For a given transcript of a protein, at exon boundaries the local context of the nucleotides was taken from the genomic context. The COSMIC consensus transcript was chosen for the transcript for each protein. Changes in codons can lead to amino acid substitutions, synonymous and nonsense mutations. Therefore, ***codon mutability*** was calculated as the probability to observe a specific type of codon change which can be realized by single nucleotide mutations at each codon position *i* as:

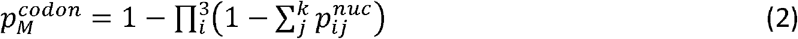

Where *k* denotes a number of mutually exclusive mutations at codon position *i*. For example, for Phe codon “TTT” in a given context 5’-A and G-3’ three single nucleotide mutations can lead to *Phe* → *Leu* substitution (to codons “TTG”, “TTA” and “CTT” for Leu): A[T≫C]TTG in the first codon position or mutually exclusive *ATT*[*T*≫*G*]*G* and *ATT*[*T*≫*A*]*G* in the third codon position. In this case the probability of *Phe* → *Leu* substitution in the *ATTTG* context can be calculated as 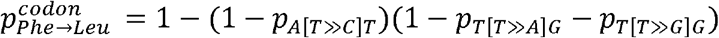 where trinucleotide frequencies were taken from the mutational profile. Amino acid substitutions corresponding to each missense mutation are calculated by translating the mutated and wild type codons using a standard codon table. Codon mutability strongly depends on the neighboring codons as illustrated in Figure S5.

### Gene-weight adjusted mutability

Gene weights estimate a relative probability of a gene compared to other genes to be mutated in cancer through somatic mutagenesis. There are multiple ways the gene weights can be calculated:

*SNP-based weight* was calculated using the number of SNPs in the vicinity of the gene of interest. We used the “EnsDb.Hsapiens.v86” database to find genomic coordinates of a gene, including introns, and extended the range in both 3’ and 5’ directions according to the window size (Table S7). We then counted the number of common SNPs from dbSNP database[51] within the genomic region. Gene weight was calculated as: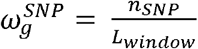, where *n_SNP_* is the number of SNPs and *L_window_* is the length of the genomic region in base pairs. We tested several window sizes for defining the genomic regions around the gene of interest (Table S6).

*Mutation-based weight* was calculated using the number of nucleotide sites with reoccurring mutations counted only once to avoid the bias that may be present due to selection on individual sites: 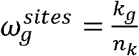. Here *k_g_* is the number of mutated sites and *n_k_* is the number of base pairs in the gene transcript.

*Silent mutation-based weight* was introduced previously and was shown to be superior in assessment of significant non-synonymous mutations across genes [52]. This weight can be calculated by taking into account only silent somatic mutations: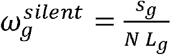. Here *s_g_* is the total number of somatic silent mutations within the gene, *N* is the number of tumor samples and *L_g_* is the number of codons in the gene transcript.

*No-outlier-based weight* introduced previously [21] takes into account the number of all codon mutations within a gene, *C_g_*, excluding mutations in outlier codon sites bearing more than the 99^th^ percentile of mutations of the gene: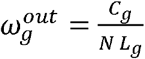, normalized by gene length *L_g_* in amino acids and the total number of samples *N*.

Using gene weights, an adjusted probability per codon can be then expressed as:

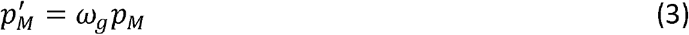

Similarly, per nucleotide probability can be calculated adjusted by gene weight.

### Identification of significant mutations

B-score uses the binomial model to calculate the probability of observing a certain type of mutation in a given site more frequently than *k*:

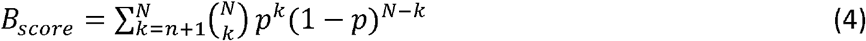

Where 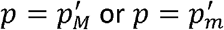 and *k* is the number of observed mutations of a given type at a particular nucleotide or codon, *N* is a total number of cancer samples in a cohort. Depending on the dataset chosen or a particular cohort of patients (for instance, corresponding to one cancer type), the total number of samples *N* and the numbers of observed mutations *k* will change. While ranking mutations in a given gene, *B_score_* can further be adjusted for multiple-testing with Benjamini-Hochberg correction as implemented on the MutaGene website.

### Computational Predictions

CanDrAplus^34^ program was downloaded and ran using default specifications with the “Cancer-in-General” annotation data file. REVEL^33^ predictions were obtained from dbNSFP database[53]. CHASMplus predictions were obtained using CRAVAT[54]. The pan-cancer model was used for CHASMplus. FatHMM^35^ cancer-associated scores were obtained from their webserver.

### Statistical analyses and evaluation of performance

Differences between various groups were tested with the Kruskal-Wallis, Dunn test, and Mann-Whitney-Wilcoxon tests implemented in *R* software. Dunn’s test is a non-parametric pairwise multiple comparisons procedure based on rank sums; it is used to infer difference between means in multiple groups and was used because it is relatively conservative post-hoc test for Kruskal-Wallis. Associations between mutability and observed frequency (the number of individuals with a mutation in whole-exome/genome studies from COSMIC), was tested using Pearson as well as Spearman correlation tests since the variables were not normally distributed.

To quantify the performance of scores, we performed Receiver Operating Characteristics (ROC) and precision-recall analyses. Sensitivity or true positive rate was defined as TPR=TP/(TP + FN) and specificity was defined as SPC=TN/(FP+TN). Additionally, in order to account for imbalances in the labeled dataset, the quality of the predictions was described by Matthews correlation coefficient:

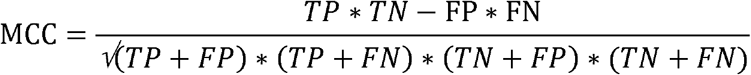

In order to compare across tools, the threshold which gave the maximum MCC was chosen for each tool to calculate TP, TN, FP, and FN.

## ACKNOWLEDGEMENT

We thank Yuri Wolf, Alejandro Schaffer and Igor Rogozin for helpful discussions. The work was supported by Intramural Research Programs of the National Library of Medicine, National Institutes of Health. Minghui Li was supported by the National Natural Science Foundation of China (Grant No. 31701136), Natural Science Foundation of Jiangsu Province, China (Grant No. BK20170335), and the Priority Academic Program Development of Jiangsu Higher Education Institutions.

---

**Figure S1.**
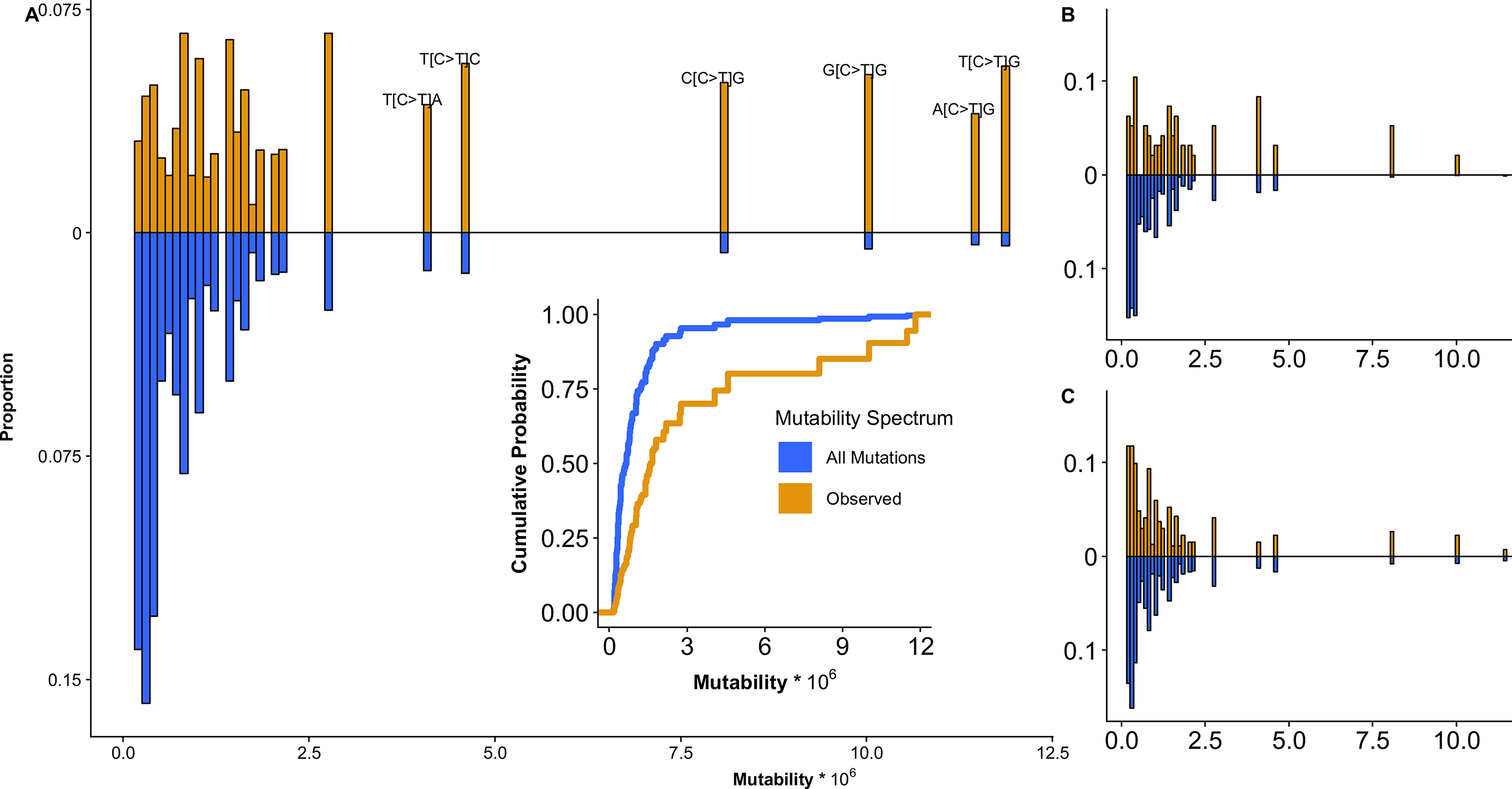
Comparison between expected nucleotide mutability spectrum of all possible mutations (blue) and mutations which were observed in cancer patients (brown) in the COSMIC v85 cohort. **(A)** Mutations from 520 cancer census genes; **(B)** CASP8 and **(C)** TP53 genes. Y-axis has been mirrored and shows the proportion of nucleotide mutations with the mutability given on the X-axis. For example, 5.6% of the 57,074 observed nucleotide mutations occurred at a site with the maximum pan-cancer nucleotide mutability of 1.18 × 10^−5^, despite the fact that only 0.4% of possible nucleotide mutations have a mutability that high. Inset shows the cumulative distribution functions for both spectra. Annotations in **(A)** show nucleotide substitutions in specific sequence contexts.

**Figure S2.**
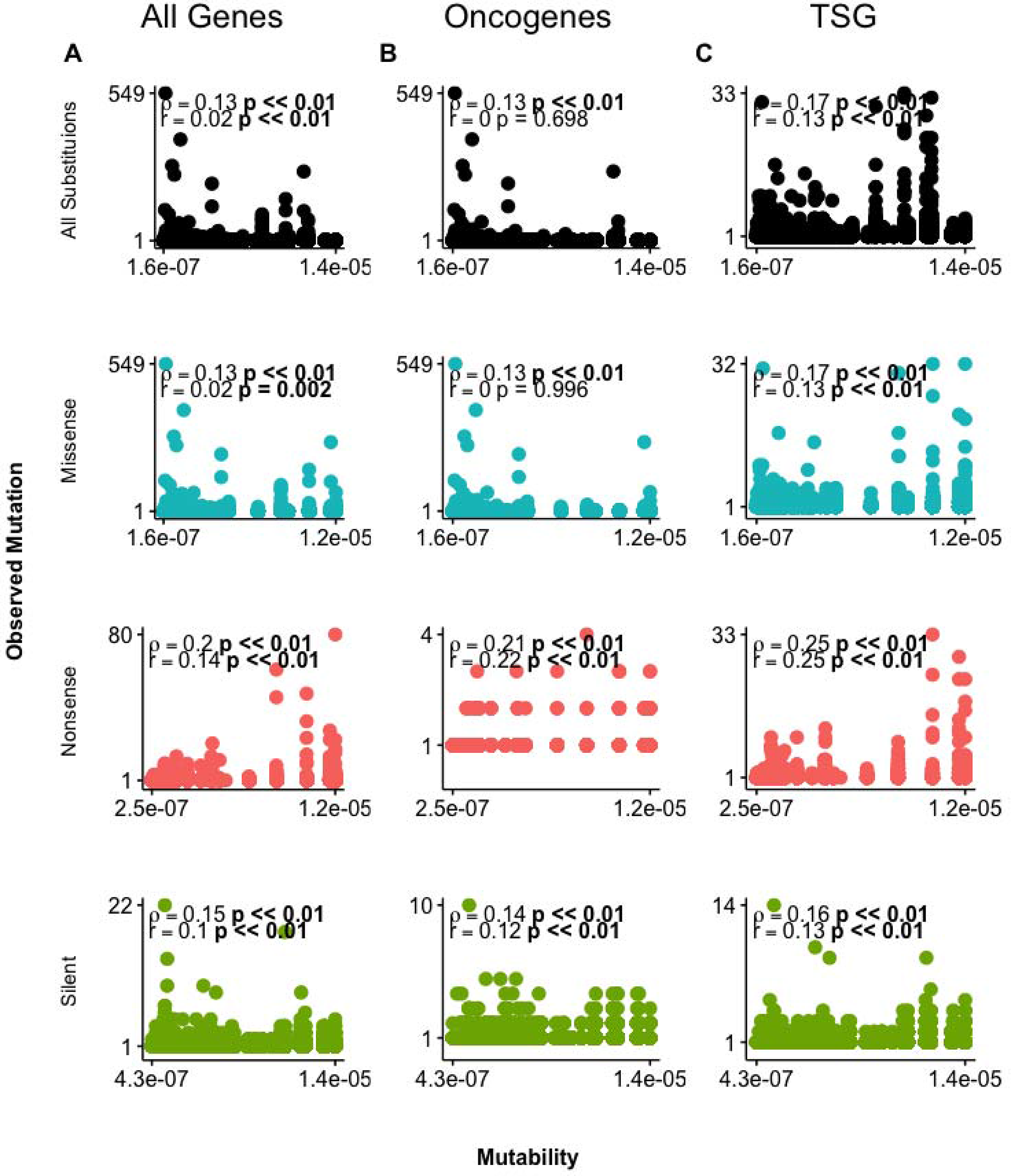
Relationship between codon mutability and reoccurrence frequency by mutation type and gene role in cancer. Scatterplots for **(A)** all cancer census genes (n = 520), **(B)** oncogenes (n = 202) and **(C)** tumor suppressor genes (TSG) (n = 166) for all mutation types: missense (blue), nonsense (red) and silent (green). Spearman and Pearson correlation coefficient with respective p-values are shown in all figures with p < 0.01 in bold.

**Figure S3.**
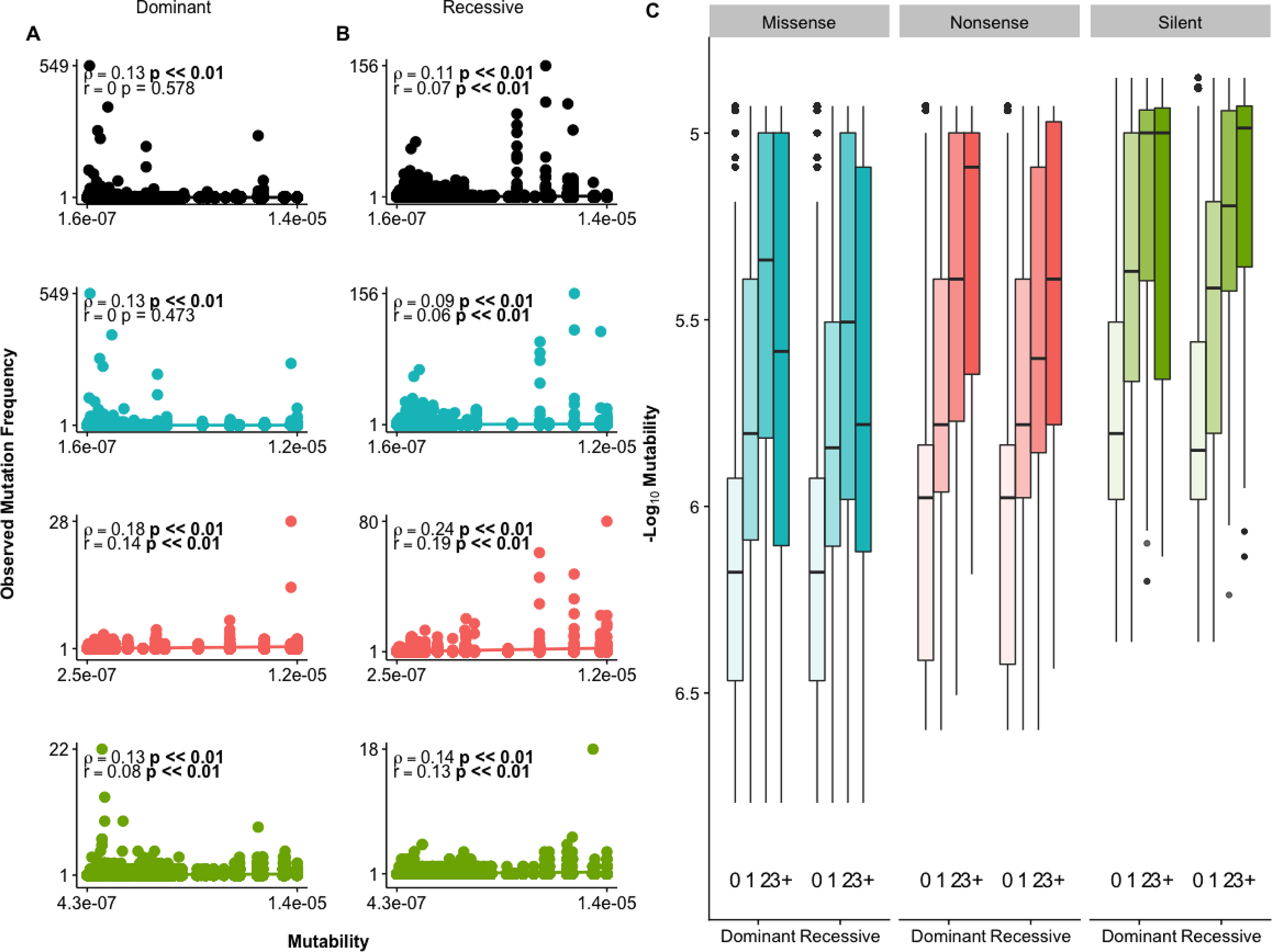
Relationship between codon mutability and observed reoccurrence frequency by mutation type and molecular genetics. **(A)** Genes with only dominant mutations, **(B)** Genes with only recessive mutations. Different colors show scatterplots broken down by mutation type: missense (blue), nonsense (red) and silent (green). **(C)** Mutations in cancer census gene grouped by Dominant and Recessive mutations. Spearman and Pearson correlation coefficient with respective p-values shown in all, significant at p < 0.01 in bold. Counts summarized in Table S1.

**Figure S4.**
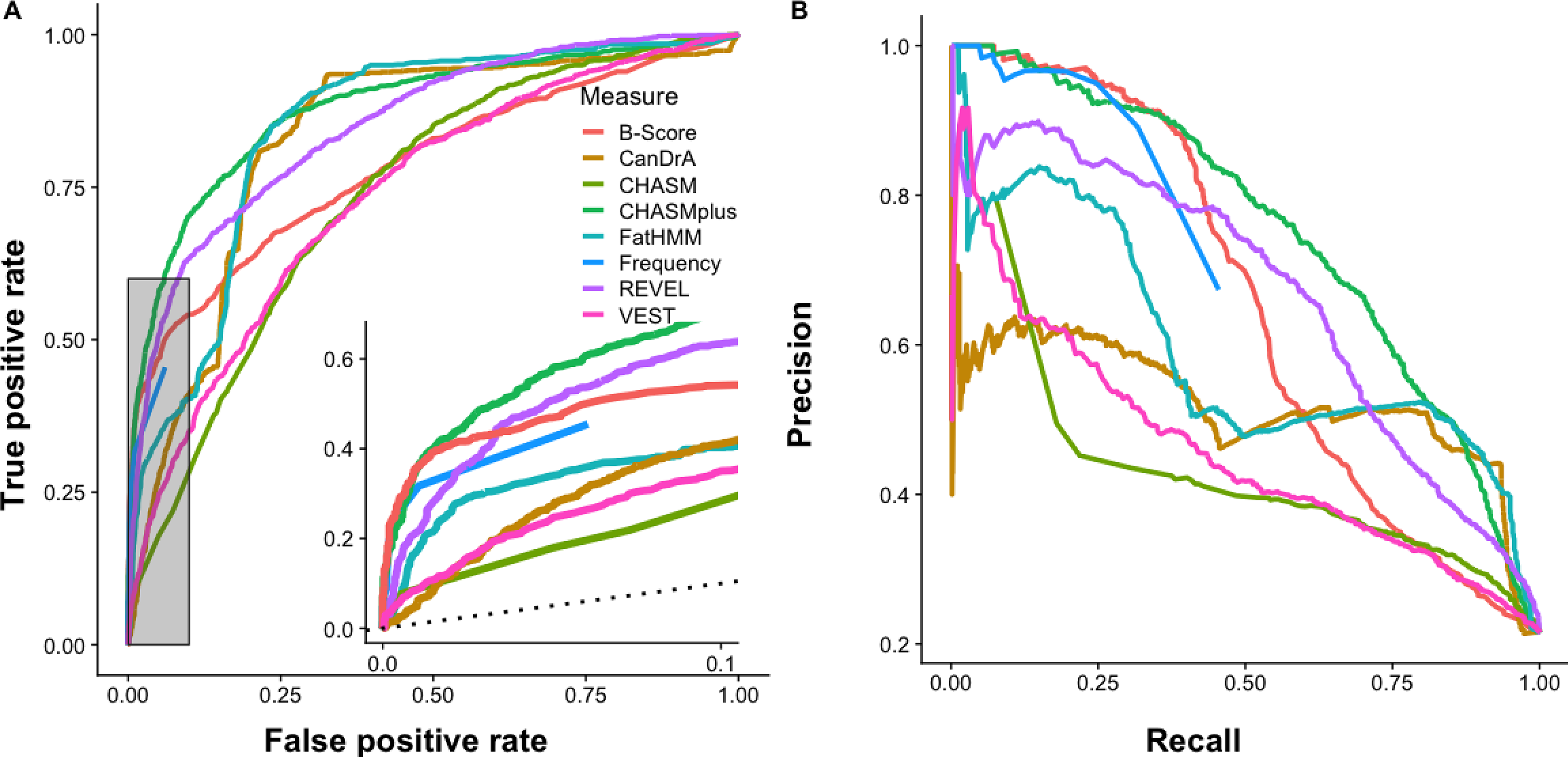
Assessment of classification performance between all neutral and non-neutral mutations in a combined dataset. **(A)** ROC curves for B-Score, and observed mutational frequency based on mutation frequency in COSMIC v85 cohort. Inset shows the performance of highlighted area corresponding to up to 10% FPR. **(B)** Precision-recall curves for the same benchmark set. The ROC for reoccurrence frequency cannot be calculated for all mutations because some experimentally validated mutations were not observed in the COSMIC v85 cohort.

**Figure S5.**
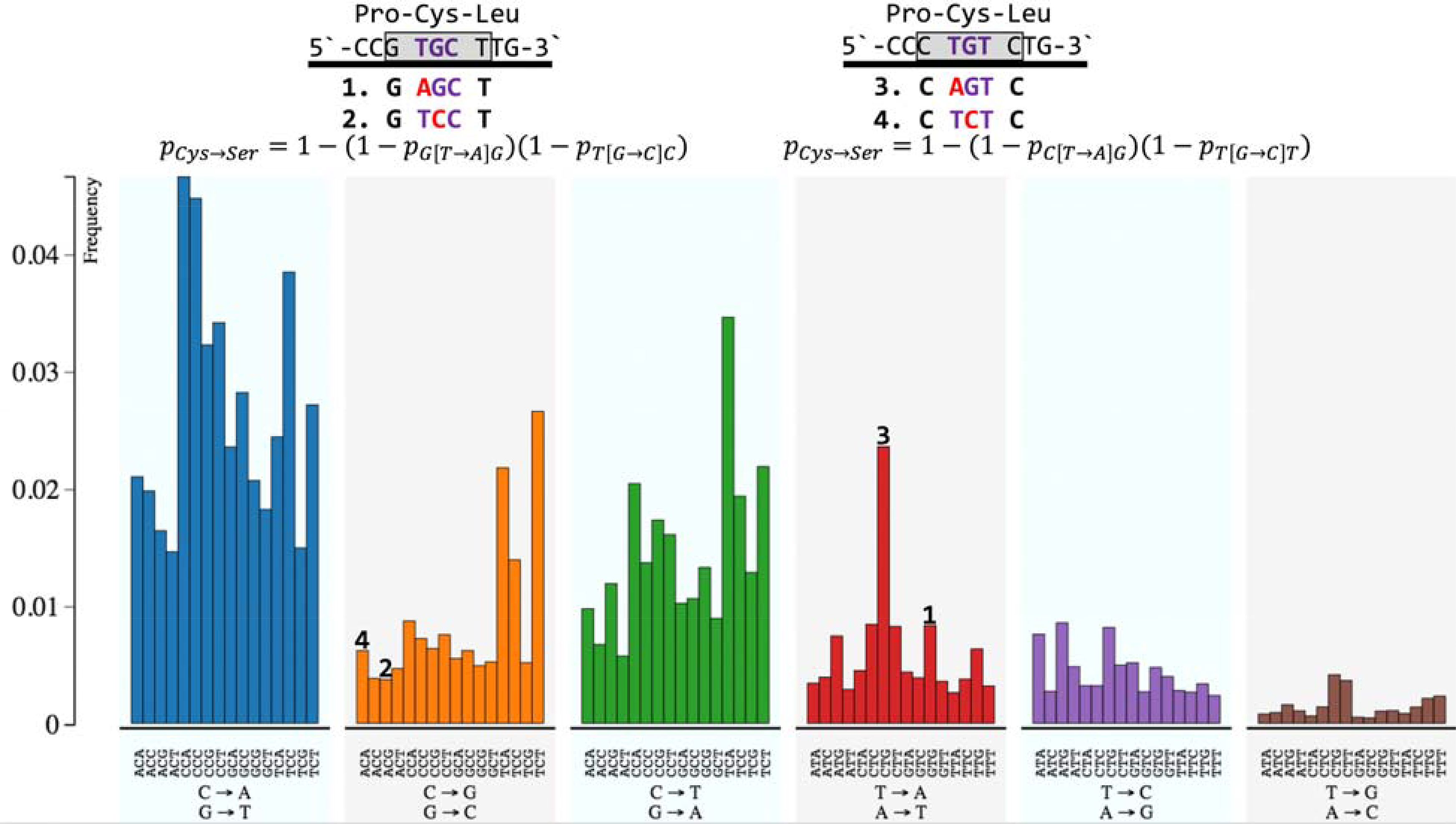
Example of how codon mutability for amino acid substitution is affected by the neighboring nucleotides. A peptide sequence of Pro-Cys-Leu could be encoded by nucleotide sequence CCG-TGC-TTG (left) or CCC-TGT-CTG (right). For both peptides, the pentanucleotides used to calculate the codon mutability for a substitution has been highlighted in the blue box. Figure below shows lung adenocarcinoma cancer mutational profile used to calculate nucleotide mutability, x-axis is the 96 different possible context-dependent mutation types, y-axis shows mutation frequency. For each of the nucleotide mutations leading to a amino acid substitution, the corresponding peak on the mutational profile id shown.

**Table S1.**
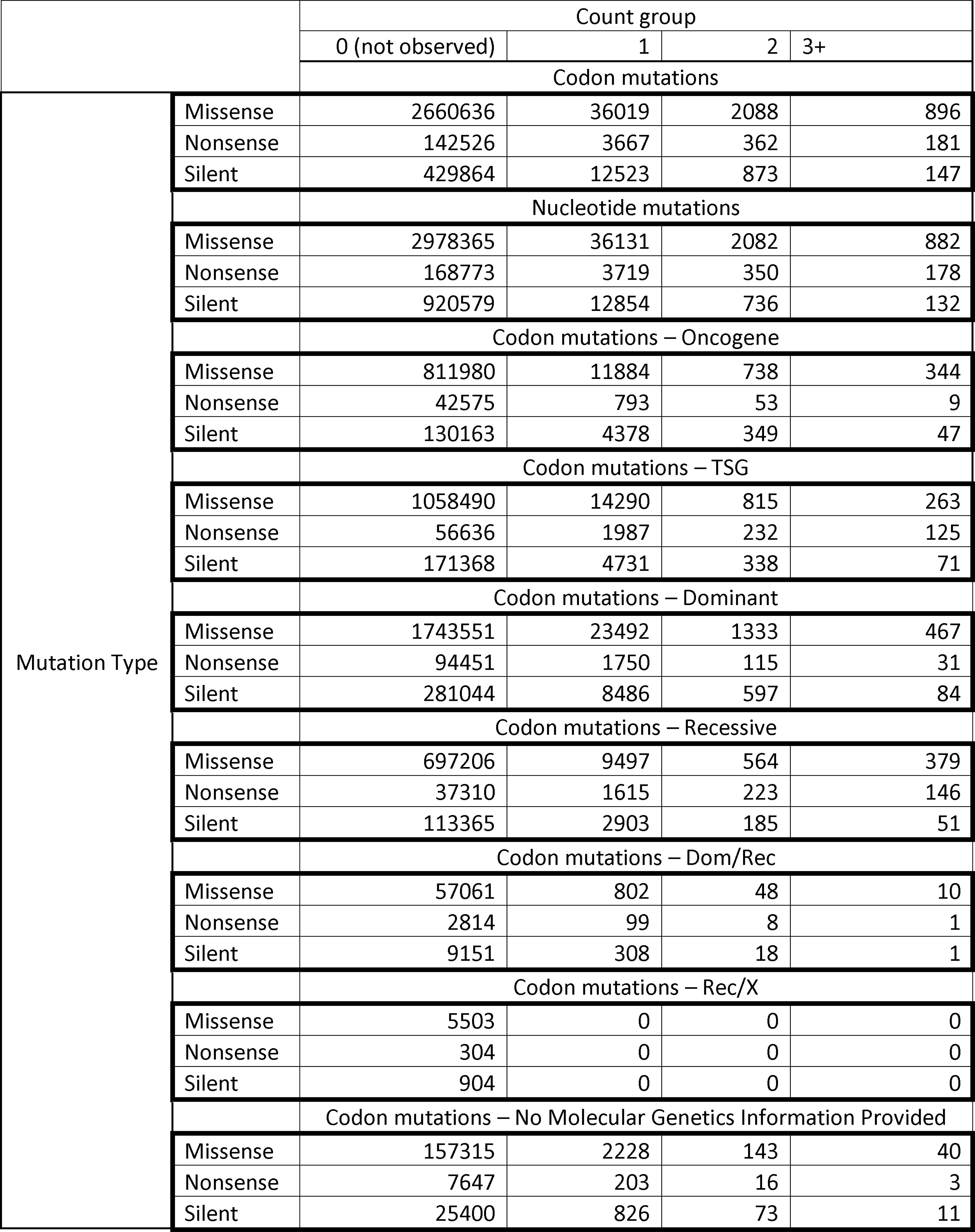
Counts for boxplots in Figure 3.

**Table S2.** Correlation between mutability and recurrence of mutations in cancer-associated genes.

https://zenodo.org/record/2575077#.XG8bA5NKjUJ

**Table S3.** List of cancer-associated genes and GO terms.

https://zenodo.org/record/2575077#.XG8bA5NKjUJ

**Table S4.** Combined dataset with experimentally annotated neutral and non-neutral mutations. in 58 genes

https://www.ncbi.nlm.ih.gov/research/mutagene/benchmark

**Table S5.**
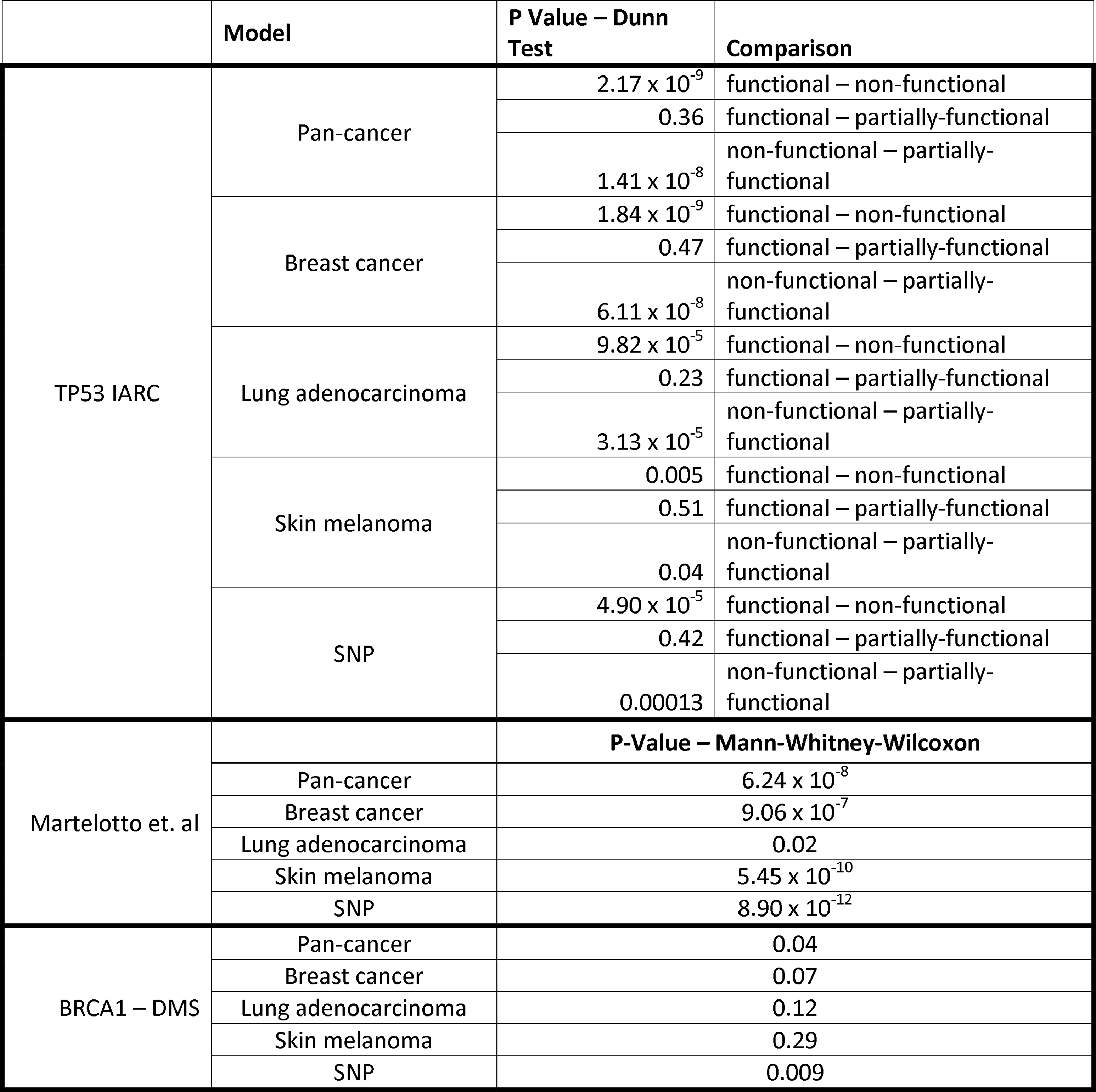
Comparison of mutability on three experimental datasets with different cancer-specific background mutation models

**Table S6.**
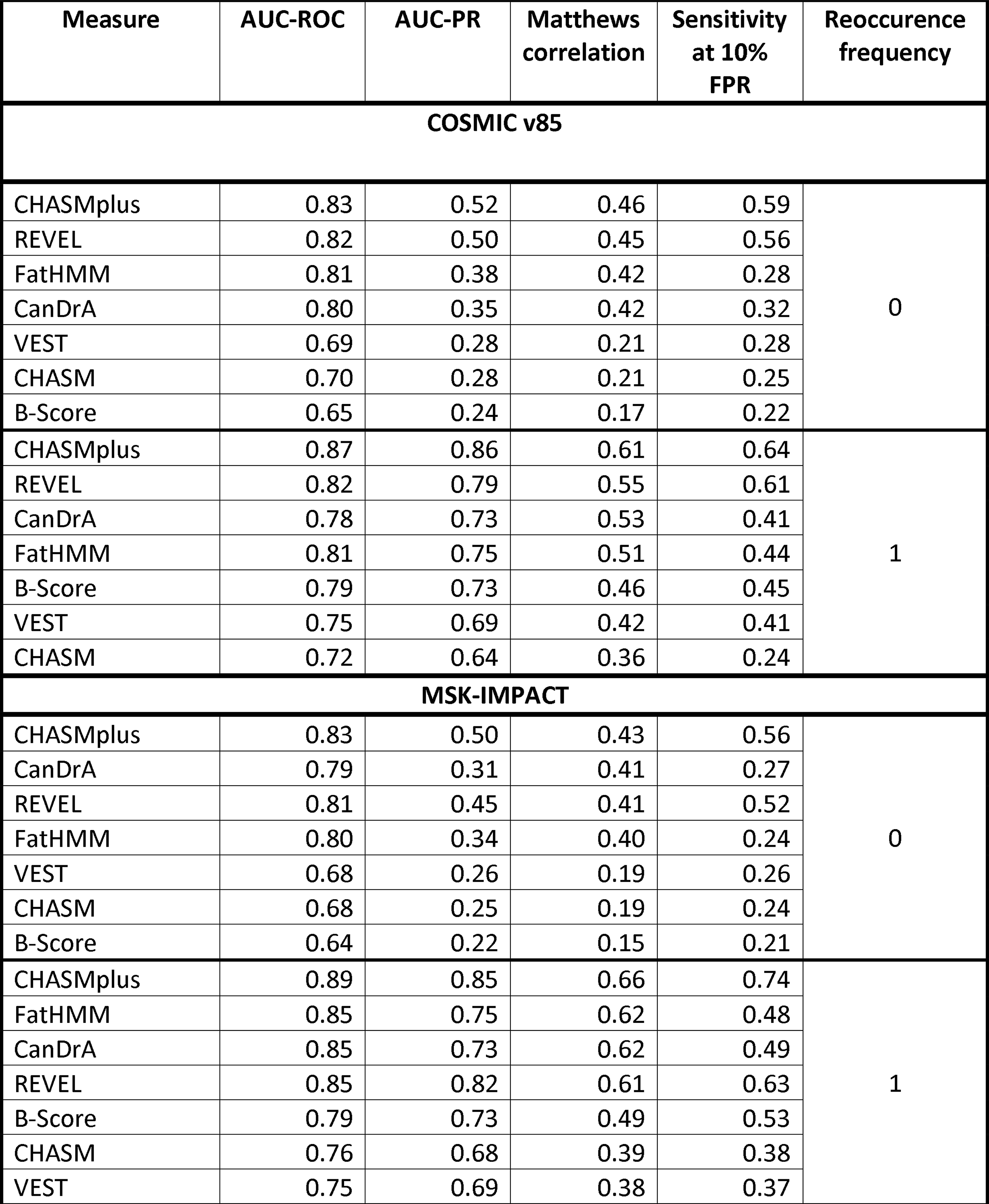
Performance of different classifiers on unobserved and rarely observed mutations in both cancer cohorts. Classifiers are sorted by their maximum Matthews correlation. B-Score for each cohort is calculated with the respective cohort size: COSMIC v85 cohort (N=12,013); MSK-Impact (N=9,228). For CHASM the background model yielding best performance was chosen.

**Table S7.**
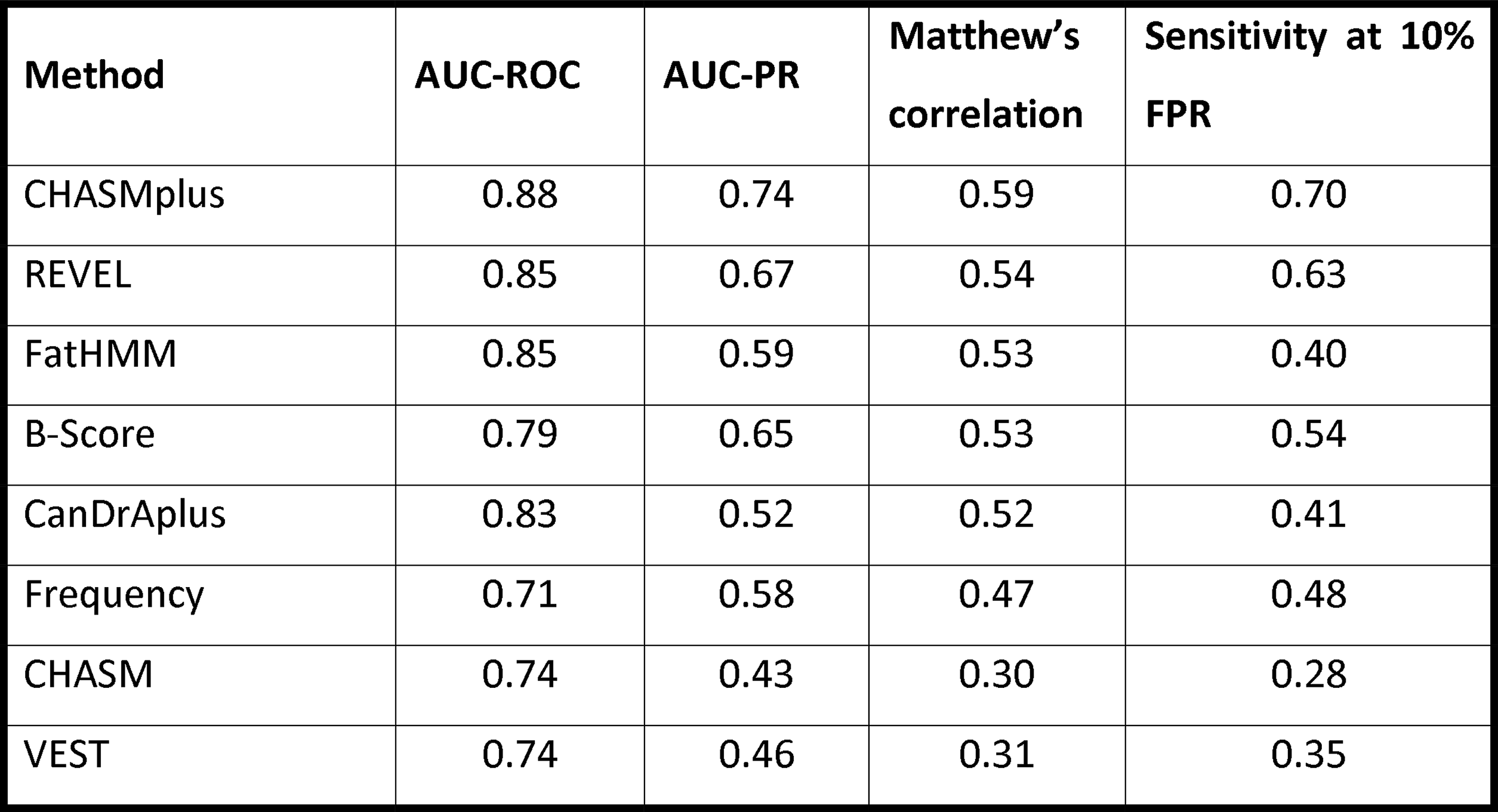
Comparison of different methods to distinguish all neutral from non-neutral mutations using *combined* experimental dataset. AUC-ROC (Area under the receiver operator curve) and AUC-PR (Area under the precision recall curve) values for reoccurrence frequency counts were extrapolated since some experimentally validated mutations were not observed in tumor samples. See Figure S6 for the ROC and PR plots. Maximum Matthew’s correlation reported for each predictor. For CHASM the background model yielding best performance was chosen.

**Table S8.**
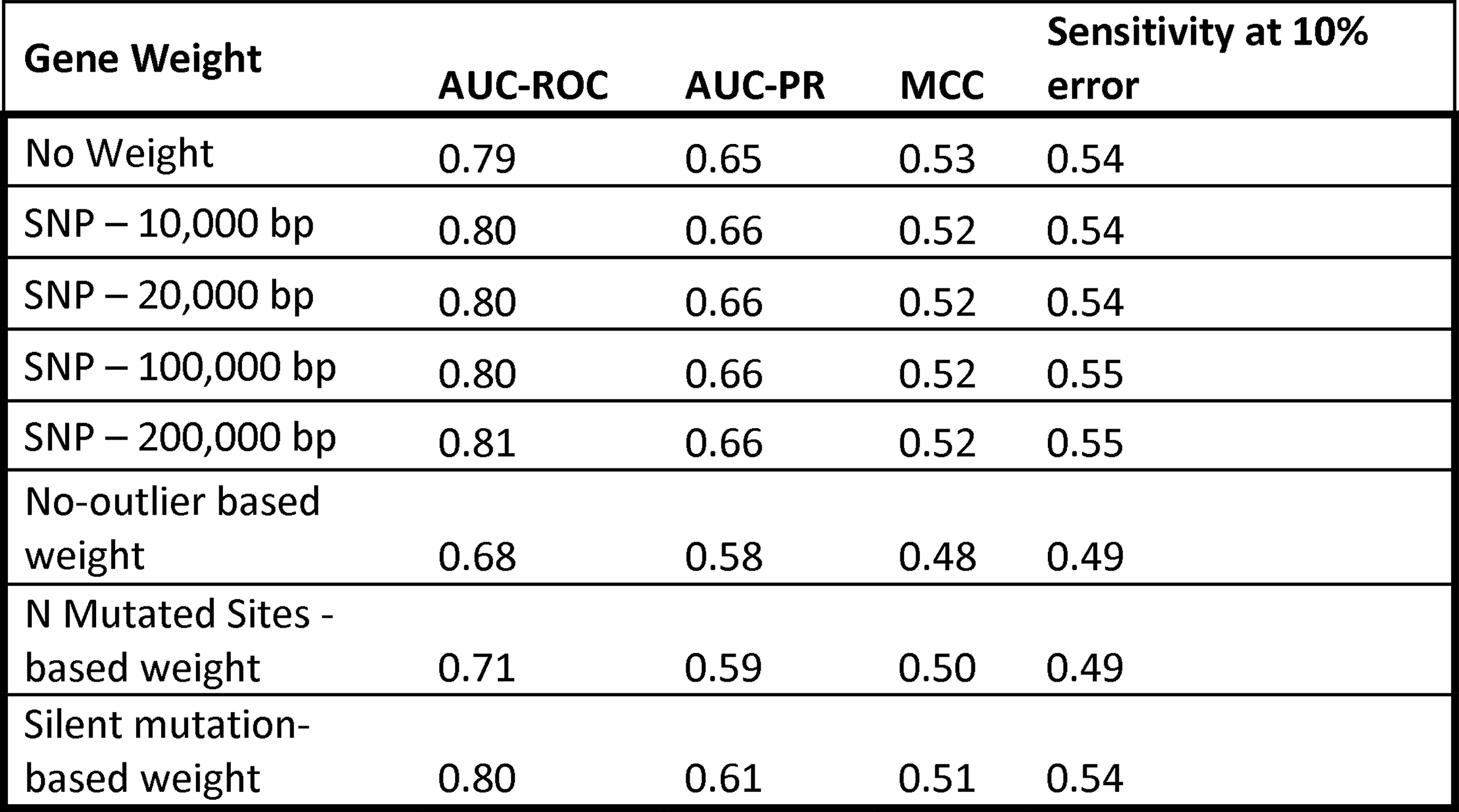
Performance metrics for B-score on *combined* dataset using different gene weights.

## References

1. Lynch M. Rate, molecular spectrum, and consequences of human mutation. Proc Natl Acad Sci U S A. 2010;107(3):961–8. Epub 2010/01/19. doi: 10.1073/pnas.0912629107. PubMed PMID: 20080596; PubMed Central PMCID: PMC2824313.

2. Greenman C, Stephens P, Smith R, Dalgliesh GL, Hunter C, Bignell G, et al. Patterns of somatic mutation in human cancer genomes. Nature. 2007;446(7132):153–8. Epub 2007/03/09. doi: 10.1038/nature05610. PubMed PMID: 17344846; PubMed Central PMCID: PMC2712719.

3. Leedham S, Tomlinson I. The continuum model of selection in human tumors: general paradigm or niche product? Cancer research. 2012;72(13):3131–4. Epub 2012/05/04. doi: 10.1158/0008-5472.CAN-12-1052. PubMed PMID: 22552286.

4. Nussinov R, Tsai CJ. ’Latent drivers’ expand the cancer mutational landscape. Curr Opin Struct Biol. 2015;32:25–32. Epub 2015/02/11. doi: 10.1016/j.sbi.2015.01.004. PubMed PMID: 25661093.

5. Dagogo-Jack I, Shaw AT. Tumour heterogeneity and resistance to cancer therapies. Nat Rev Clin Oncol. 2018;15(2):81–94. Epub 2017/11/09. doi: 10.1038/nrclinonc.2017.166. PubMed PMID: 29115304. 6.

6. Hiley C, de Bruin EC, McGranahan N, Swanton C. Deciphering intratumor heterogeneity and temporal acquisition of driver events to refine precision medicine. Genome Biol. 2014;15(8):453. Epub 2014/09/16. doi: 10.1186/s13059-014-0453-8. PubMed PMID: 25222836; PubMed Central PMCID: PMCPMC4281956.

7. Nik-Zainal S, Alexandrov LB, Wedge DC, Van Loo P, Greenman CD, Raine K, et al. Mutational processes molding the genomes of 21 breast cancers. Cell. 2012;149(5):979–93. Epub 2012/05/23. doi: 10.1016/j.cell.2012.04.024. PubMed PMID: 22608084; PubMed Central PMCID: PMCPMC3414841.

8. Alexandrov LB, Nik-Zainal S, Wedge DC, Aparicio SA, Behjati S, Biankin AV, et al. Signatures of mutational processes in human cancer. Nature. 2013;500(7463):415–21. Epub 2013/08/16. doi: 10.1038/nature12477. PubMed PMID: 23945592; PubMed Central PMCID: PMC3776390.

9. Rogozin IB, Pavlov YI, Goncearenco A, De S, Lada AG, Poliakov E, et al. Mutational signatures and mutable motifs in cancer genomes. Briefings in bioinformatics. 2017. Epub 2017/05/13. doi: 10.1093/bib/bbx049. PubMed PMID: 28498882.

10. Michaelson JJ, Shi Y, Gujral M, Zheng H, Malhotra D, Jin X, et al. Whole-genome sequencing in autism identifies hot spots for de novo germline mutation. Cell. 2012;151(7):1431–42. Epub 2012/12/25. doi: 10.1016/j.cell.2012.11.019. PubMed PMID: 23260136; PubMed Central PMCID: PMC3712641.

11. Hodgkinson A, Eyre-Walker A. Variation in the mutation rate across mammalian genomes. Nature reviews Genetics. 2011;12(11):756–66. Epub 2011/10/05. doi: 10.1038/nrg3098. PubMed PMID: 21969038.

12. Lynch M, Ackerman MS, Gout JF, Long H, Sung W, Thomas WK, et al. Genetic drift, selection and the evolution of the mutation rate. Nature reviews Genetics. 2016;17(11):704–14. Epub 2016/10/16. doi: 10.1038/nrg.2016.104. PubMed PMID: 27739533.

13. Stratton MR. Exploring the genomes of cancer cells: progress and promise. Science. 2011;331(6024):1553–8. Epub 2011/03/26. doi: 10.1126/science.1204040. PubMed PMID: 21436442.

14. Polak P, Karlic R, Koren A, Thurman R, Sandstrom R, Lawrence M, et al. Cell-of-origin chromatin organization shapes the mutational landscape of cancer. Nature. 2015;518(7539):360–4. Epub 2015/02/20. doi: 10.1038/nature14221. PubMed PMID: 25693567; PubMed Central PMCID: PMC4405175.

15. Rogozin IB, Kolchanov NA. Somatic hypermutagenesis in immunoglobulin genes. II. Influence of neighbouring base sequences on mutagenesis. Biochimica et biophysica acta. 1992;1171(1):11–8. Epub 1992/11/15. PubMed PMID: 1420357.

16. Chen C, Qi H, Shen Y, Pickrell J, Przeworski M. Contrasting Determinants of Mutation Rates in Germline and Soma. Genetics. 2017;207(1):255–67. Epub 2017/07/25. doi: 10.1534/genetics.117.1114. PubMed PMID: 28733365; PubMed Central PMCID: PMC5586376.

17. Gonzalez-Perez A, Lopez-Bigas N. Functional impact bias reveals cancer drivers. Nucleic acids research. 2012;40(21):e169. Epub 2012/08/21. doi: 10.1093/nar/gks743. PubMed PMID: 22904074; PubMed Central PMCID: PMC3505979.

18. Martincorena I, Raine KM, Gerstung M, Dawson KJ, Haase K, Van Loo P, et al. Universal Patterns of Selection in Cancer and Somatic Tissues. Cell. 2017;171(5):1029–41 e21. Epub 2017/10/24. doi: 10.1016/j.cell.2017.09.042. PubMed PMID: 29056346; PubMed Central PMCID: PMC5720395.

19. Araya CL, Cenik C, Reuter JA, Kiss G, Pande VS, Snyder MP, et al. Identification of significantly mutated regions across cancer types highlights a rich landscape of functional molecular alterations. Nature genetics. 2016;48(2):117–25. Epub 2015/12/23. doi: 10.1038/ng.3471. PubMed PMID: 26691984; PubMed Central PMCID: PMC4731297.

20. Peterson TA, Gauran IIM, Park J, Park D, Kann MG. Oncodomains: A protein domain-centric framework for analyzing rare variants in tumor samples. PLoS computational biology. 2017;13(4):e1005428. Epub 2017/04/21. doi: 10.1371/journal.pcbi.1005428. PubMed PMID: 28426665; PubMed Central PMCID: PMC5398485.

21. Chang MT, Asthana S, Gao SP, Lee BH, Chapman JS, Kandoth C, et al. Identifying recurrent mutations in cancer reveals widespread lineage diversity and mutational specificity. Nature biotechnology. 2016;34(2):155–63. Epub 2015/12/01. doi: 10.1038/nbt.3391. PubMed PMID: 26619011; PubMed Central PMCID: PMC4744099.

22. Porta-Pardo E, Kamburov A, Tamborero D, Pons T, Grases D, Valencia A, et al. Comparison of algorithms for the detection of cancer drivers at subgene resolution. Nature methods. 2017;14(8):782–8. Epub 2017/07/18. doi: 10.1038/nmeth.4364. PubMed PMID: 28714987.

23. Wong WC, Kim D, Carter H, Diekhans M, Ryan MC, Karchin R. CHASM and SNVBox: toolkit for detecting biologically important single nucleotide mutations in cancer. Bioinformatics. 2011;27(15):2147–8. Epub 2011/06/21. doi: 10.1093/bioinformatics/btr357. PubMed PMID: 21685053; PubMed Central PMCID: PMC3137226.

24. Li M, Kales SC, Ma K, Shoemaker BA, Crespo-Barreto J, Cangelosi AL, et al. Balancing Protein Stability and Activity in Cancer: A New Approach for Identifying Driver Mutations Affecting CBL Ubiquitin Ligase Activation. Cancer research. 2016;76(3):561–71. Epub 2015/12/18. doi: 10.1158/0008-5472.CAN-14-3812. PubMed PMID: 26676746; PubMed Central PMCID: PMCPMC4738050.

25. Campbell BB, Light N, Fabrizio D, Zatzman M, Fuligni F, de Borja R, et al. Comprehensive Analysis of Hypermutation in Human Cancer. Cell. 2017;171(5):1042–56 e10. Epub 2017/10/24. doi: 10.1016/j.cell.2017.09.048. PubMed PMID: 29056344; PubMed Central PMCID: PMCPMC5849393.

26. Bailey MH, Tokheim C, Porta-Pardo E, Sengupta S, Bertrand D, Weerasinghe A, et al. Comprehensive Characterization of Cancer Driver Genes and Mutations. Cell. 2018;173(2):371–85 e18. Epub 2018/04/07. doi: 10.1016/j.cell.2018.02.060. PubMed PMID: 29625053.

27. Molina-Vila MA, Nabau-Moreto N, Tornador C, Sabnis AJ, Rosell R, Estivill X, et al. Activating mutations cluster in the “molecular brake” regions of protein kinases and do not associate with conserved or catalytic residues. Hum Mutat. 2014;35(3):318–28. Epub 2013/12/11. doi: 10.1002/humu.22493. PubMed PMID: 24323975.

28. Schaafsma GCP, Vihinen M. Large differences in proportions of harmful and benign amino acid substitutions between proteins and diseases. Hum Mutat. 2017;38(7):839–48. Epub 2017/04/27. doi: 10.1002/humu.23236. PubMed PMID: 28444810.

29. Stehr H, Jang SH, Duarte JM, Wierling C, Lehrach H, Lappe M, et al. The structural impact of cancer-associated missense mutations in oncogenes and tumor suppressors. Mol Cancer. 2011;10:54. Epub 2011/05/18. doi: 10.1186/1476-4598-10-54. PubMed PMID: 21575214; PubMed Central PMCID: PMCPMC3123651.

30. Chang MT, Bhattarai TS, Schram AM, Bielski CM, Donoghue MTA, Jonsson P, et al. Accelerating Discovery of Functional Mutant Alleles in Cancer. Cancer Discov. 2018;8(2):174–83. Epub 2017/12/17. doi: 10.1158/2159-8290.CD-17-0321. PubMed PMID: 29247016; PubMed Central PMCID: PMCPMC5809279.

31. Carter H, Chen S, Isik L, Tyekucheva S, Velculescu VE, Kinzler KW, et al. Cancer-specific high-throughput annotation of somatic mutations: computational prediction of driver missense mutations. Cancer research. 2009;69(16):6660–7. Epub 2009/08/06. doi: 10.1158/0008-5472.CAN-09-1133. PubMed PMID: 19654296; PubMed Central PMCID: PMCPMC2763410.

32. Tokheim C, Karchin R. CHASMplus reveals the scope of somatic missense mutations driving human cancers. 2019. doi: https://doi.org/10.1101/313296.

33. Douville C, Masica DL, Stenson PD, Cooper DN, Gygax DM, Kim R, et al. Assessing the Pathogenicity of Insertion and Deletion Variants with the Variant Effect Scoring Tool (VEST-Indel). Hum Mutat. 2016;37(1):28–35. Epub 2015/10/08. doi: 10.1002/humu.22911. PubMed PMID: 26442818; PubMed Central PMCID: PMCPMC5057310.

34. Ioannidis NM, Rothstein JH, Pejaver V, Middha S, McDonnell SK, Baheti S, et al. REVEL: An Ensemble Method for Predicting the Pathogenicity of Rare Missense Variants. Am J Hum Genet. 2016;99(4):877–85. Epub 2016/09/27. doi: 10.1016/j.ajhg.2016.08.016. PubMed PMID: 27666373; PubMed Central PMCID: PMCPMC5065685.

35. Mao Y, Chen H, Liang H, Meric-Bernstam F, Mills GB, Chen K. CanDrA: cancer-specific driver missense mutation annotation with optimized features. PLoS One. 2013;8(10):e77945. Epub 2013/11/10. doi: 10.1371/journal.pone.0077945. PubMed PMID: 24205039; PubMed Central PMCID: PMCPMC3813554.

36. Shihab HA, Gough J, Cooper DN, Stenson PD, Barker GL, Edwards KJ, et al. Predicting the functional, molecular, and phenotypic consequences of amino acid substitutions using hidden Markov models. Hum Mutat. 2013;34(1):57–65. Epub 2012/10/04. doi: 10.1002/humu.22225. PubMed PMID: 23033316; PubMed Central PMCID: PMC3558800.

37. Lawrence MS, Stojanov P, Polak P, Kryukov GV, Cibulskis K, Sivachenko A, et al. Mutational heterogeneity in cancer and the search for new cancer-associated genes. Nature. 2013;499(7457):214–8. Epub 2013/06/19. doi: 10.1038/nature12213. PubMed PMID: 23770567; PubMed Central PMCID: PMCPMC3919509.

38. Goncearenco A, Rager SL, Li M, Sang QX, Rogozin IB, Panchenko AR. Exploring background mutational processes to decipher cancer genetic heterogeneity. Nucleic acids research. 2017;45(W1):W514–W22. Epub 2017/05/05. doi: 10.1093/nar/gkx367. PubMed PMID: 28472504; PubMed Central PMCID: PMCPMC5793731.

39. Gorlov IP, Gorlova OY, Amos CI. Relative effects of mutability and selection on single nucleotide polymorphisms in transcribed regions of the human genome. BMC Genomics. 2008;9:292. Epub 2008/06/19. doi: 10.1186/1471-2164-9-292. PubMed PMID: 18559102; PubMed Central PMCID: PMCPMC2442617.

40. Supek F, Minana B, Valcarcel J, Gabaldon T, Lehner B. Synonymous mutations frequently act as driver mutations in human cancers. Cell. 2014;156(6):1324–35. Epub 2014/03/19. doi: 10.1016/j.cell.2014.01.051. PubMed PMID: 24630730.

41. Ainscough BJ, Griffith M, Coffman AC, Wagner AH, Kunisaki J, Choudhary MN, et al. DoCM: a database of curated mutations in cancer. Nat Methods. 2016;13(10):806–7. Epub 2016/09/30. doi: 10.1038/nmeth.4000. PubMed PMID: 27684579; PubMed Central PMCID: PMCPMC5317181.

42. Landrum MJ, Lee JM, Riley GR, Jang W, Rubinstein WS, Church DM, et al. ClinVar: public archive of relationships among sequence variation and human phenotype. Nucleic Acids Res. 2014;42(Database issue):D980–5. Epub 2013/11/16. doi: 10.1093/nar/gkt1113. PubMed PMID: 24234437; PubMed Central PMCID: PMCPMC3965032.

43. Olivier M, Eeles R, Hollstein M, Khan MA, Harris CC, Hainaut P. The IARC TP53 database: new online mutation analysis and recommendations to users. Hum Mutat. 2002;19(6):607–14. Epub 2002/05/15. doi: 10.1002/humu.10081. PubMed PMID: 12007217.

44. Martelotto LG, Ng CK, De Filippo MR, Zhang Y, Piscuoglio S, Lim RS, et al. Benchmarking mutation effect prediction algorithms using functionally validated cancer-related missense mutations. Genome Biol. 2014;15(10):484. Epub 2014/10/29. doi: 10.1186/s13059-014-0484-1. PubMed PMID: 25348012; PubMed Central PMCID: PMC4232638.

45. Starita LM, Young DL, Islam M, Kitzman JO, Gullingsrud J, Hause RJ, et al. Massively Parallel Functional Analysis of BRCA1 RING Domain Variants. Genetics. 2015;200(2):413–22. Epub 2015/04/01. doi: 10.1534/genetics.115.175802. PubMed PMID: 25823446; PubMed Central PMCID: PMC4492368.

46. Mahmood K, Jung CH, Philip G, Georgeson P, Chung J, Pope BJ, et al. Variant effect prediction tools assessed using independent, functional assay-based datasets: implications for discovery and diagnostics. Hum Genomics. 2017;11(1):10. Epub 2017/05/18. doi: 10.1186/s40246-017-0104-8. PubMed PMID: 28511696; PubMed Central PMCID: PMC5433009.

47. Ng PK, Li J, Jeong KJ, Shao S, Chen H, Tsang YH, et al. Systematic Functional Annotation of Somatic Mutations in Cancer. Cancer Cell. 2018;33(3):450–62 e10. Epub 2018/03/14. doi: 10.1016/j.ccell.2018.01.021. PubMed PMID: 29533785; PubMed Central PMCID: PMCPMC5926201.

48. Forbes SA, Beare D, Boutselakis H, Bamford S, Bindal N, Tate J, et al. COSMIC: somatic cancer genetics at high-resolution. Nucleic Acids Res. 2017;45(D1):D777–D83. Epub 2016/12/03. doi: 10.1093/nar/gkw1121. PubMed PMID: 27899578; PubMed Central PMCID: PMC5210583.

49. Gao J, Aksoy BA, Dogrusoz U, Dresdner G, Gross B, Sumer SO, et al. Integrative analysis of complex cancer genomics and clinical profiles using the cBioPortal. Sci Signal. 2013;6(269):pl1. Epub 2013/04/04. doi: 10.1126/scisignal.2004088. PubMed PMID: 23550210; PubMed Central PMCID: PMCPMC4160307.

50. Barretina J, Caponigro G, Stransky N, Venkatesan K, Margolin AA, Kim S, et al. The Cancer Cell Line Encyclopedia enables predictive modelling of anticancer drug sensitivity. Nature. 2012;483(7391):603–7. Epub 2012/03/31. doi: 10.1038/nature11003. PubMed PMID: 22460905; PubMed Central PMCID: PMCPMC3320027.

51. Sherry ST, Ward MH, Kholodov M, Baker J, Phan L, Smigielski EM, et al. dbSNP: the NCBI database of genetic variation. Nucleic Acids Res. 2001;29(1):308–11. Epub 2000/01/11. PubMed PMID: 11125122; PubMed Central PMCID: PMC29783.

52. Evans P, Avey S, Kong Y, Krauthammer M. Adjusting for background mutation frequency biases improves the identification of cancer driver genes. IEEE Trans Nanobioscience. 2013;12(3):150–7. Epub 2013/05/23. doi: 10.1109/TNB.2013.2263391. PubMed PMID: 23694700; PubMed Central PMCID: PMC3989533.

53. Liu X, Wu C, Li C, Boerwinkle E. dbNSFP v3.0: A One-Stop Database of Functional Predictions and Annotations for Human Nonsynonymous and Splice-Site SNVs. Hum Mutat. 2016;37(3):235–41. Epub 2015/11/12. doi: 10.1002/humu.22932. PubMed PMID: 26555599; PubMed Central PMCID: PMCPMC4752381.

54. Douville C, Carter H, Kim R, Niknafs N, Diekhans M, Stenson PD, et al. CRAVAT: cancer-related analysis of variants toolkit. Bioinformatics. 2013;29(5):647–8. Epub 2013/01/18. doi: 10.1093/bioinformatics/btt017. PubMed PMID: 23325621; PubMed Central PMCID: PMCPMC3582272.

